# Satellite DNA dynamics across phylogenetic scales in ground beetles and other insects

**DOI:** 10.64898/2026.05.27.728298

**Authors:** Zachary S. Warner, Sherif Negm, Patrick Wynn, Tuan Pham, Lorraine Zaki, Gilbert Giri, Paul B. Frandsen, Amanda M. Larracuente, John S. Sproul

**Affiliations:** Brigham Young University, Department of Biology, Provo, UT; University of Rochester, Department of Biology, Rochester, NY; University of Chicago, Department of Human Genetics, IL; Brown University, Center for Computational Molecular Biology, Data Science Institute, Providence, RI; Touro College of Osteopathic Medicine, Middletown, NY; University of North Carolina School of Medicine, Bioinformatics and Computational Biology, Chapel Hill, NC; Brigham Young University, Department of Plant and Wildlife Science, Provo, UT; University of Nebraska Omaha, Department of Biology, Omaha, NE

**Keywords:** tandem repeats, satDNA, comparative genomics, rapid genome evolution, Insecta, Carabidae, *Bembidion*

## Abstract

Satellite DNAs (satDNAs) are tandemly repeated sequences that can comprise large fractions of eukaryotic genomes. For decades, we have known that satellite DNAs are among the most rapidly evolving genomic components, yet few studies have examined their dynamics across broad evolutionary frameworks—likely in part because their rapid evolution complicates or precludes homology identification. We investigated satDNA dynamics across 50 ground beetle (Carabidae) species, sampling above and below the species level in the subgenus *Plataphus* of *Bembidion*. We used RepeatExplorer2 and a novel homology detection pipeline, which we validated using species of *Drosophila* for which satDNAs are well-documented, to study satDNA dynamics in a comparative framework. We show that satDNAs comprise large genomic fractions (mean = 25.7%, range = 3.2%–53%), with individual satDNA motifs accounting for >30% of total DNA within some species. We quantify major satDNA turnover among closely related species which drives extensive repatterning of repeat content over short evolutionary scales. To place our findings in a broader context, we also surveyed satDNA diversity across the beetle suborder (Adephaga), the beetle order (Coleoptera), and the next six largest insect orders (Lepidoptera, Diptera, Hymenoptera, Hemiptera, Orthoptera, Trichoptera), representing 400 additional species. While satDNA abundance was significantly elevated in the focal subgenus (*Plataphus*), high genomic proportions were also found in species across Adephaga, Coleoptera, Hymenoptera, and Diptera. Conversely, satDNA proportions were notably reduced in Lepidoptera and Trichoptera. Our investigation of satDNA dynamics at narrow and broad taxonomic scales provides a quantitative illustration of satDNA turnover across a well-sampled clade. It also identifies new high-potential systems for gaining mechanistic insights into the drivers and impacts of satDNA evolution, and raises questions regarding potential links between satDNA turnover, speciation, and core strategies of chromosome and genome architecture in diverse clades.

## Introduction

Satellite DNAs (satDNAs) are tandemly repeated sequence motifs that can comprise large fractions of eukaryotic genomes [1,2]. In many organisms, satDNAs accumulate in sequence blocks that can span megabases of chromosomes, particularly near areas of reduced recombination such as centromeres and telomeres [3,4].

The roles satDNAs play in genomic and evolutionary processes are a point of ongoing investigation [5–8]. They can play critical roles in chromatin formation and chromosome architecture [9–12], as regulators of gene expression [13,14], and as key players in meiotic drive systems [4,15–17]. Their rapid evolution can be a major source of genomic differentiation among species [18–21] and has been linked to hybrid incompatibilities and the process of speciation [22–24].

Despite their ubiquity and known impacts on genomic and evolutionary processes, satDNAs are understudied relative to other DNA sequence types (e.g., genic, regulatory, and transposable elements). This may be partly because satDNAs can complicate molecular biology experiments and genomics research, thus making some insights technically challenging or intractable [25–27]. In addition, the rapid nature of satDNA evolution makes individual satDNAs somewhat ephemeral over evolutionary time [3]. Unlike genes and transposable elements which contain functionally constrained sequences (e.g., open reading frames), satDNAs frequently lack such constraints and may thus evolve rapidly at the sequence level [28]. This means that the phylogenetic distribution of their sequences may be limited to an individual species or their close relatives. The ephemeral nature of satDNA creates a barrier of entry when approaching a new study system, because extensive *de novo* characterization of satDNAs will likely be required.

Barriers to entry are stronger still when conducting comparative analysis across species. Although conservation of individual satDNAs is not expected across deep evolutionary time, an area of lagging understanding is how tandem repeats like satDNAs broadly shape genome structure and content across lineages at broad phylogenetic scales.

Despite challenges associated with studying satDNAs, approaching satDNA evolution from a many-species evolutionary framework is a potentially powerful lens for deepening understanding of satDNA dynamics and identifying ideal model systems. Most studies examine satDNA in the context of one or a few species [29–34]. A smaller number of studies have examined satDNA dynamics across broader phylogenetic scales [35–39], but these analyses are typically limited to small clades or conduct relatively sparse taxon sampling relative to species diversity.

In this study, we investigated satDNA dynamics across 50 ground beetle species within a single subgenus with dense sampling both above and below the species level. To address technical challenges associated with comparative analysis across species, we developed a computational pipeline for automated homology detection of satDNAs. We extended our investigation by characterizing satDNA dynamics in 400 insect species across progressively broader phylogenetic scales, expanding from our focal subgenus to the beetle suborder Adephaga, then to Coleoptera (i.e., all beetles), and finally to six other species-rich insect orders.

## Methods

### Taxonomic sampling

We investigated satDNA dynamics in 50 ground beetle species densely sampled from within the subgenus *Plataphus* of *Bembidion* (Carabidae). These samples were taken from data sets published previously by the last author [40,41]. *Plataphus* is a well-supported subgenus of *Bembidion* [42] and is thought to have diversified mostly in the last 10–20 million years [43]. The sampling included here targeted multiple species from every major clade within the subgenus and includes all known species from some clades [41]. For one such group, a complex of nine recently delimited species called the *breve* species group, we sampled 31 individuals across eight of the nine species, with replicate specimens sampled from multiple geographic localities. Thus, our sampling in the target subgenus spans 81 specimens sampled both above and below the species level. We downloaded shotgun Illumina data for each sample from Sequence Read Archive (SRA) BioProjects PRJNA369077 and PRJNA596448 [40,41]. Prior to downstream analysis we removed adapters and quality-trimmed raw reads using TrimGalore v0.6.3 [44]. In addition to sampling in the subgenus *Plataphus*, we sampled 400 species belonging to other beetle and insect groups described in more detail below, and in Supplemental Methods.

Supplemental file S1 provides a summary of our taxonomic sampling and SRA accessions.

### satDNA identification

We identified candidate satDNA sequences by analyzing trimmed Illumina reads in RepeatExplorer2 [45] using the TAREAN tool [46]. We subsampled reads to ∼0.5X coverage prior to analysis by calculating average coverage of 10 single-copy genes in RepeatProfiler v1.1 [47] and downsampling reads using seqtk (available from https://github.com/lh3/seqtk). We considered as candidate satDNAs those sequences identified by TAREAN as high-and low-confidence satellites.

### SatDNA homology detection pipeline

A number of tools exist for *de novo* identification of satDNA sequences both from raw reads and genome assemblies [6]. For evolutionary studies in groups without a prior history of satDNA work, it is usually necessary to conduct *de novo* homology detection of predicted satDNA sequences across samples. However, this can be challenging for multiple reasons. First, unlike many genomic regions (e.g., protein-coding genes, transposable elements, regulatory regions), satDNAs lack structural landmarks (e.g., start/stop codons, open reading frames, regulatory sequences) that define start and end points or approximate sequence length. Because of this, any sequences present in the raw output of satDNA identification software programs are not in any standardized frame of reference across samples. In addition, because satDNAs often lack evolutionary constraints at the sequence level, they can evolve rapidly both in terms of sequence and length. Thus, decisions about homology cutoffs will produce inconsistent outcomes across satDNA sequences without deliberate standardization. In order to more efficiently identify putatively homologous satDNAs across study species, we developed a satDNA homology inference pipeline in the early stages of the research.

Our approach takes as input a FASTA file containing a list of satDNA sequences for each sample. In our case, this was a file containing putative satDNAs predicted by TAREAN for all samples. The pipeline then uses each satDNA sequence as a BLAST query against a custom BLAST library containing satDNA sequences from all other samples. Next, potential homologs are identified as those which are reciprocal best hits that pass both length and sequence similarity filters. The initial sequence length and similarity filters we used cluster any sequences that share at least 90% sequence similarity and least 80% length similarity. Prior to applying the length and similarity filters, it is necessary to standardize start and endpoints of potential homologs. This standardization proceeds by aligning all sequences within a preliminary satDNA homology group to a dimerized version of that homolog (generated by selecting the longest sequence in the group and dimerizing that sequence). By aligning to a dimerized repeat, potential homologs from different species have a chance to align well, even if the satDNA detection software (i.e., TAREAN) set different start and end points for that sequence in each species. A random position in that alignment is then chosen at which sequences to the left are cut and added to the sequence on the right followed by realignment to the dimerized reference. In this way, start and endpoints are standardized, and length and sequence similarity filters can be applied without alignment artefacts due to variable definitions in monomer start/end points across species. Any sequences that remain grouped following these filters are accepted as putative homologs and a consensus sequence of each is generated. The pipeline then outputs a list of satDNA homology groups and a FASTA file of consensus sequences for each group.

We validated our satDNA detection methods and homology detection pipeline using six *Drosophila* species with well-documented satDNAs: *D. melanogaster* (SRA accession: SRR6399448), *D. melanogaster* (SRR6399448)*, D. simulans* (SRR6399446)*, D. sechellia* (SRR6399452)*, D. mauritiana* (SRR6425993), *D. erecta* (SRR6425990), and *D. yakuba* (SRR6426004). We trimmed, downsampled reads, and identified satDNAs with RepeatExplorer2 using the same methods as the beetles reported above. We then ran the satDNAs detected by RepeatExplorer2 through our homology detection pipeline. We predicted that if our combined use of RepeatExplorer2 output with our homology pipeline was generating reliable data, the results would correctly identify abundant satDNAs known to occur across species, and reflect known patterns of presence/absence of those satDNAs as follows: *Rsp* (*D. melanogaster* only), *Rsp-like* (*D. simulans*, *D. mauritiana,* and *D. sechellia*) [48], and the *1.688* family satDNAs (*D. melanogaster*, *D. simulans*, *D. mauritiana, D. sechellia, D. yakuba, and D. erecta*) [49].

### Quantification and visualization of satDNA profiles

In addition to initial analyses in RepeatExplorer2, we quantified satDNA abundance and visualized satDNA profiles using RepeatProfiler [47] with two general approaches.

RepeatProfiler maps reads to a suite of reference sequences and outputs visually enhanced read depth profiles in addition to a run summary table with read mapping results for each reference sequence. In our first approach, we used RepeatProfiler to map reads from each species to the full suite of satDNAs identified within the same respective species. We analyzed RepeatProfiler run summary tables to calculate the overall genomic abundance of satDNA, the genomic proportion and copy number of each individual satDNA after normalization relative to single-copy protein-coding genes, and the number of satDNAs which account for 50% and 90% of reads mapped to the total suite of satDNAs, respectively. We plotted the resulting data using custom scripts in R v4.0.2 [50].

Our second approach focused on comparing satDNA profiles at finer evolutionary scales, among and within species in the *breve* species group. For this approach, we mapped reads from all samples of *breve* group species to the same satDNA library which contained only one representative sequence (i.e., from a single species) for each satDNA identified within the *breve* group species. This approach of mapping reads from multiple samples to the same library of satDNAs derived from a single representative species can allow for comparative study of profiles with a standard frame of reference when study species show minimal evolutionary divergence [47]. For satellites that were shared among multiple species, the species with the longest consensus sequence was chosen to represent that satDNA. In all RepeatProfiler runs, we analyzed 5 million read pairs for each sample. Although we were primarily interested in raw read mapping values, we also converted raw coverage depths to copy number estimates. We normalized read depth relative to 10 single-copy genes from *Bembidion lividulum*, which we added to the file containing satDNA reference sequences and using the “-single_copy” flag in our RepeatProfiler command. We visualized the resulting patterns with graphical output generated by RepeatProfiler which we subsequently arranged in Adobe Illustrator.

### Characterizing satellite DNA turnover across *Plataphus* species

We used four complementary approaches to investigate satDNA turnover across species. First, we visualized satDNA abundance metrics in their evolutionary context alongside a phylogeny of *Plataphus* to qualitatively evaluate whether closely related species showed major shifts in overall satDNA abundance, or in the abundance of specific satDNA sequences. Second, we applied formal tests of phylogenetic signal to our data using Blomberg’s *K* [51] and Pagel’s λ [52]. We used these tests to statistically evaluate whether patterns of (1) satDNA genomic proportion, (2) count of total satDNAs, and (3) the genomic proportion of the single most abundant satDNA across species are strongly predicted by phylogenetic relatedness. We reasoned that strong phylogenetic signal in satDNA metrics would indicate modest rates of satDNA turnover, whereas low levels of phylogenetic signal would indicate relatively high rates of turnover. We assessed statistical significance of Blomberg’s *K* using a permutation test with 1000 replicates and assessed significance against a null hypothesis of Brownian motion. We assessed statistical significance for Pagel’s λ using a likelihood ratio test. All tests were conducted using the APE v5.8.1 [53] and phytools v2.5.2 [54] R packages in R v4.5.3 [50] with a custom R script. Third, we evaluated the extent to which homologous satellite DNAs detected by our homology analysis described above were conserved across species. Fourth, we used results of read mapping to satDNA references generated by RepeatProfiler [47] as described above to quantify shifts in abundance of specific satDNAs on a species-by-species basis and visualized patterns as a heatmap generated using a custom script in R.

### Validating short-read estimates with long-read data in *Bembidion laxatum*

Following observations of striking individual satDNA motif abundance in *B. laxatum* (described in more detail below), we sought a second estimate of satDNA abundance using long-read data for that species. High molecular weight DNA extraction was conducted on fresh tissue of a single male specimen at the Center for Genomics (CGB) and Bioinformatics at Brigham Young University. The CGB prepared a HiFi sequencing library using the SMRT bell prep kit 3.0 following standard protocols. The library was sequenced on the PacBio Revio instrument. We assembled the HiFi reads using Hifiasm v.0.2.5 [55]. We removed haplotypic variation from the assembly using purge_dups with cutoffs set from manual inspection of the kmer histogram. We assessed gene completeness of the assembly using compleasm v.0.2.6 [56] with the Endopterygota odb10 BUSCO set. We identified and classified repetitive elements in the assembly using RepeatModeler2 [57] and RepeatMasker [58] as implemented in the EarlGrey v7.0.1 pipeline [59]. We then converted both the curated repeat library produced by EarlGrey and the *B. laxatum* assembly into custom BLAST databases using Geneious Prime v2023.2.1 [60]. We then conducted BLAST searches using as queries the consensus sequences of satDNAs generated by RepeatExplorer2 as part of our short-read approach described above.

This approach allowed us to evaluate both the strikingly high estimates of a specific satDNA sequence and more generally validate the satDNA consensus sequences produced by RepeatExplorer2 using an orthogonal approach.

### Survey of satDNA dynamics across major insect orders

To compare satDNA dynamics observed in *Plataphus* to broader evolutionary scales, we expanded our analysis to include 400 additional insect species including species form the beetle suborder Adephaga, the beetle order Coleoptera, and the six next most species-rich insect orders (Diptera, Hemiptera, Hymenoptera, Lepidoptera, Orthoptera, and Trichoptera). We briefly summarize here our methods for standardizing and filtering our final dataset with additional details provided in Supplemental Methods. We retrieved publicly available Illumina shotgun data (paired-end 150 bp, Illumina, whole-genome – see Supplemental Methods for more criteria used to control variables expected in public data) from the Sequence Read Archive (SRA), randomly selecting species within each order with a limit of one species per genus. To ensure representative taxonomic breadth, we verified representation of major families within each order using the Catalogue of Life Checklist [61], leading to the inclusion of five additional samples from the family Curculionidae to address a sampling gap of a major lineage in Coleoptera. Our sampling of adephagan species did not include any species within subgenus *Plataphus* (or even the genus *Bembidion*) and therefore represents adephagan estimates independent of *Plataphus*.

We estimated satDNA abundance using RepeatExplorer2’s TAREAN module [46]. Because most species lack published genome-size estimates, we identified the smallest chromosome-scale assembly within each order on NCBI to estimate a lower bound of genome size. Reads for all species in an order were then downsampled to the 1x equivalent of this minimum genome size. While this conservative strategy ensured all samples likely conformed to the ≤1x coverage requirement for TAREAN, it resulted in lower coverage (e.g., 1/10 coverage) for species with larger genomes, potentially reducing sensitivity for low abundance satDNAs. However, we preferred this loss in sensitivity of low abundance satDNAs rather than risk violating assumptions of the analysis with samples that might exceed 1x coverage.

We performed post-analysis filtering by visualizing summaries of RepeatExplorer2 output for all samples. Samples for which over 90% of reads were assigned to repeat clusters were further investigated, leading to the removal of 22 runs derived from museum specimens with putatively low-quality DNA, or non-focal sources. One high-abundance outlier, *Mylabris sibirica* (68.2% genomic proportion of satDNA), was retained only after validation via BLAST against an independent long-read assembly for that species [62] (see Supplemental Methods for more validation details). To standardize sample sizes, we used a Python script that evaluated 50,000 random subsets of 50 samples for each lineage, then kept the subsets that most closely reflected the original satDNA proportion for that respective lineage. Finally, we used R v4.5.3 [50] to visualize patterns of satDNA dynamics across focal groups. Following analysis of the full dataset, we conducted a sensitivity analysis to evaluate the impacts of limited sampling on the mean values within orders. We calculated order-wide trends using a dataset that randomly sampled half the number of species and compared those results with those of our fully sampled dataset.

## Results

### Satellite DNA abundance

Satellite DNAs comprise large genomic fractions of nearly all *Plataphus* species with an average genomic proportion of 25.7% with 11 of 50 species showing ≥ 40%, two of which are >50% (Fig. 1). SatDNA abundance accounts for an average of 51.0% (stdev = 11.8) of all repetitive sequences in *Plataphus* (Figs. 1 and 2). The total number of satDNAs detected by RepeatExplorer2 within a given species ranges from 7 to 45 with an average of 25.5 (stdev = 9.3). Total satDNA counts are not correlated with the genomic proportion of satDNA, or overall genomic repetitiveness (Fig. 2C, D). In most species, a small number of the total satDNA sequences account for most of the overall satDNA abundance. We quantified this by mapping all genomic reads to the full suite of satDNAs detected by RepeatExplorer2 and then asking which satellites accounted for the majority of the total reads mapped. On average 2.4 satellites account for 50% of reads mapped to the total suite of satDNAs, and 7.4 satellites account for 90% of mapped reads (Figs. 1, 2B). The single most abundant satellite comprises an average genomic proportion of 8.9% (stdev = 5.9%) but exceeds 20% in three species (*B. laxatum*, *B. rusticum lenensoides*, *B. gordoni*) (Fig. 1).

**Figure 1.**
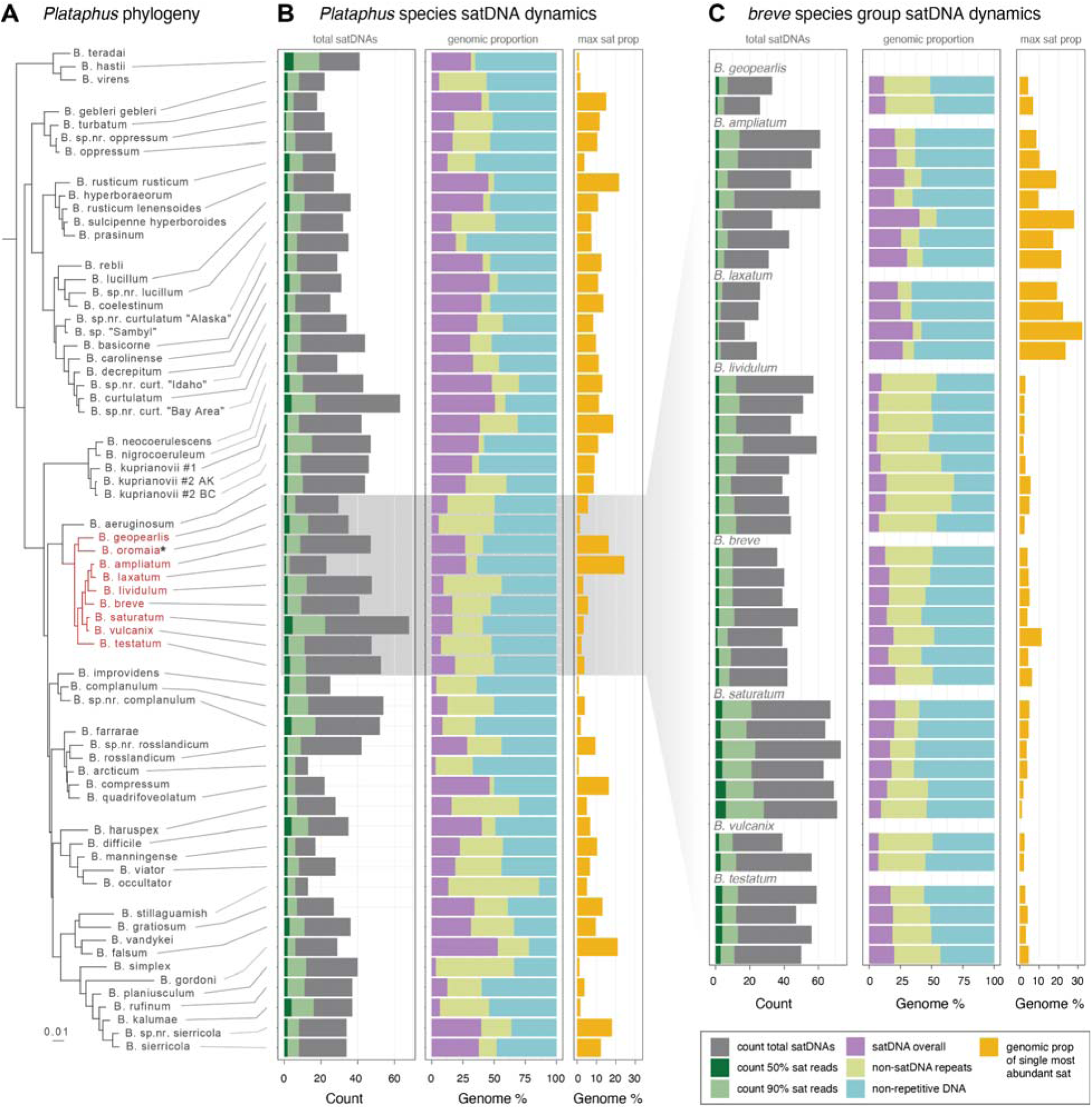
SatDNA dynamics in *Plataphus* and the *breve* species group. (A) Phylogeny of Plataphus showing the relationships as they were presented in Sproul and Maddison [41]. The *breve* group species for which sampling within species was conducted and shown in B are shown in red text. (B) Summary of satDNA dynamics across ground beetle species in the subgenus *Plataphus* of *Bembidion* (Carabidae). Values for species belonging to the *breve* species group shown are within-species averages calculated from the specimen-by-specimen numbers reported in C. (C) Summary of satDNA dynamics with multiple replicates replicate specimens sampled within species of the *breve* species group of *Plataphus*. For both B and C, the first respective column summarizes overall counts of satDNAs (gray), the count of satDNAs that account for 90% of reads mapping to the full suite of satDNAs (light green), and the count of satDNAs that account for 50% of all reads mapping to the full suite of satDNAs (dark green). The second respective column summarizes the genomic proportions of satDNA (purple), repetitive DNA overall (yellow), and non-repetitive sequences (teal). The third respective column summarizes the genomic proportion of the single most abundant satDNA detected. \**Bembidion oromaia* is represented in panel B, however due to a lack of replicate specimens for that species it is omitted from Panel C despite being a species in the *breve* species group.

**Figure 2.**
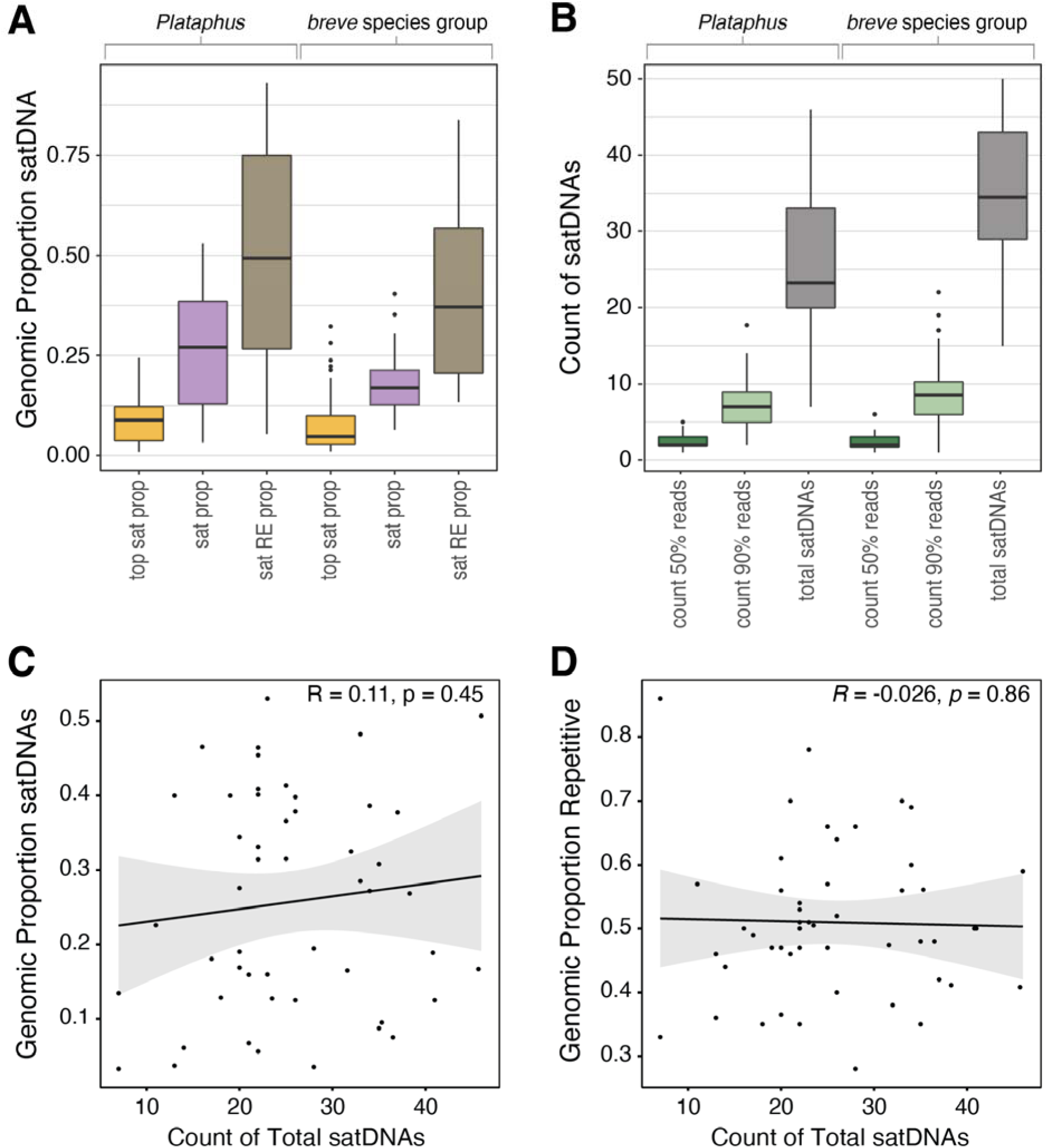
SatDNA summary statistics and correlations. (A) Genomic proportion of satDNA for each of three categories: “top sat prop” shows the genomic proportion of the single most abundant satDNA motif, “sat prop” shows the genomic proportion of all satDNAs detected, and “sat RE prop” shows the proportion of all repetitive elements that is comprised of satDNA. Colors correspond to categories shown in Figure 1 (except that “sat RE prop” is now represented in Figure 1). (B) Summarizes counts of satDNA motifs: “sat count 50% reads” shows how many satDNA motifs account for 50% of all reads mapping to satDNA, “sat count 90% reads” shows how many satDNA motifs account for 90% of all reads mapping to satDNA, and “total satDNAs” shows how many total satDNA motifs were detected. Colors correspond to categories shown in Figure 1. (C) Scatterplot showing the relationship between genomic proportion of satDNAs and the count of total satDNAs within *Plataphus* species expressed as Pearson correlation coefficient. (D) Scatterplot showing the relationship between genomic proportion of repetitive sequences and the count of total satDNAs within *Plataphus* species expressed as Pearson correlation coefficient.

At finer taxonomic scales in the *breve* species group of *Plataphus* we sampled multiple individuals per species. SatDNA abundance shows generally modest within-species variation across replicate specimens from different geographic localities (Fig. 1B). We quantified variation within vs between species using a variance partitioning analysis which examined three measures of satDNA dynamics: genomic proportion of satDNA, the genomic proportion of the single most abundant satDNA motif, and the satDNA proportion of total repeats. This analysis showed strong species-level signal in satDNA variation across the three metrics analyzed.

Comparisons between species accounted for 4.2x, 5.6x, and 20.8x more of the variance than within-species comparisons for genomic proportion of satDNA, the genomic proportion of the single most abundant satDNA motif, and the satDNA proportion of total repeats, respectively. Intraclass correlation coefficients-like values were 0.80, 0.84, and 0.95 for the same respective metrics. Taken together, these variance partition analysis results indicate that differences between species account for the large majority of overall variation in the data.

### Satellite DNA turnover and homology detection

Mapping satDNA dynamics to the *Plataphus* phylogeny showed many cases in which satDNA counts, overall genomic proportion, and abundance of the single most common satDNA motif show major shifts among sister taxa and closely related species (Fig. 1). These observations were corroborated by formal tests of phylogenetic signal. The effects of phylogeny had weak explanatory power in accounting for satDNA genomic proportion across species. Blomberg’s *K* was less than 1 (*K = 0.36,* P = 0.024), suggesting within-clade similarity is lower than expected. The significant p-value suggests more phylogenetic structure than random but taken together these values suggest overall weak phylogenetic signal in satDNA abundance. Pagel’s λ was a non-zero and non-significant value (λ = 0.264, P = 0.318) suggesting the data are statistically consistent with no phylogenetic signal (Fig. S1). The same tests showed mixed results when testing for phylogenetic signal of total satDNA counts (i.e., the count of different satDNA motifs) within species indicating limited but modest phylogenetic signal (*K* = 0.329, P = 0.067; λ = 0.484, P = 0.022) (Fig. S2). Finally, the genomic proportion of the single most abundant satellite showed no evidence of phylogenetic signal with a non-significant *K* value much less than one (*K* = 0.297, P = 0.113) and a non-significant λ near 0 (λ = >0.001, P = 1) (Fig. S3). Together, these phylogenetic signal tests provide further evidence of high overall rates of satDNA turnover on relatively short timescales in *Plataphus*.

Our homology pipeline automated detection of putatively homologous satDNAs across study species and helped overcome technical challenges associated with multi-species satDNA alignment (Fig. 3). Homology analysis within *Plataphus* showed 163 homology groups in which a putatively homologous satDNA was detected in at least two different species. On average each homology group contained 3.30 species (stdev = 2.70) and only seven of the 163 homology groups included ten or more species. In cases where several species were present within homology groups, the group composition was typically limited to closely related species, often sister taxa, or species belonging to the same subclade/species group (Fig. 4). We also observed many cases in which satDNAs that comprised >10% genomic proportion in a species were either absent or showed very low abundance in their sister taxon (e.g., see Homology Group 20 in *Bembidion neoceorulescens*, Homology Group 63 in *B*. *turbatum,* and Homology Group 68 in *B. rusticum lenensoides*) (Fig. 4). The low number of species per homology group, the limited number of homology groups containing broad species membership, and the presence of many cases of major variation in satDNA abundance between closely related species demonstrate how dynamic turnover of satDNAs account for wholesale change of large genomic fractions across species in this subgenus.

**Figure 3.**
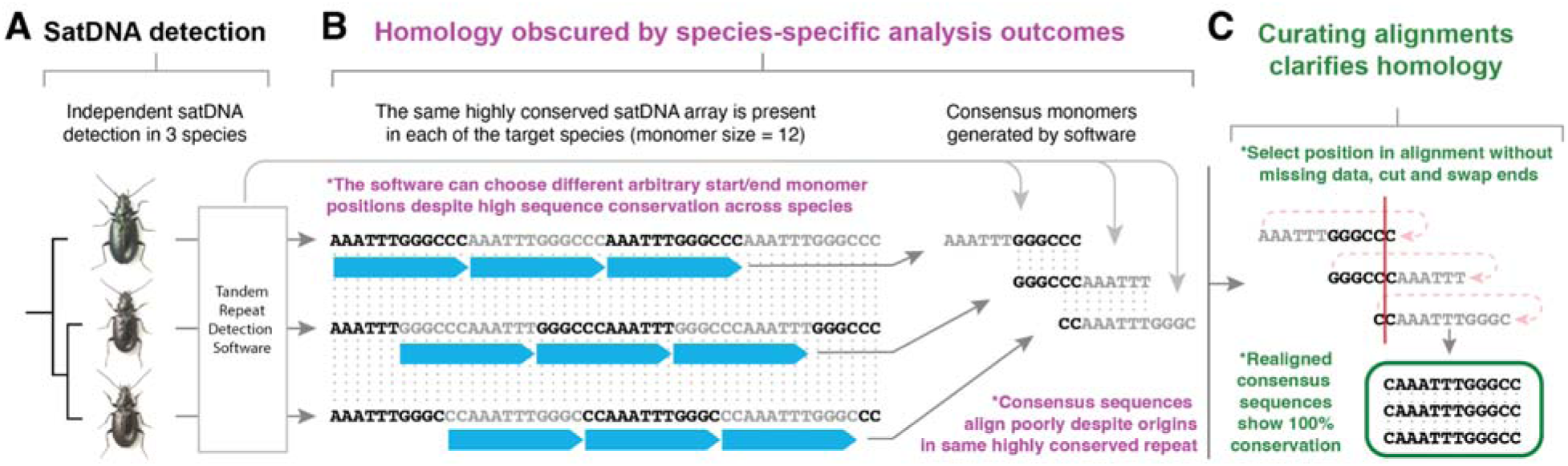
Technical challenge and solution to improve satDNA homology detection across species. Detecting homology in tandemly repeated sequences can be challenging even when closely related species share highly conserved satDNAs. (A) Three species are independently analyzed with satDNA detection software which will identify satDNAs, define monomers and output consensus sequences of detected satDNAs. (B) Because species are analyzed independently, software algorithms may define monomer boundaries in different ways across species. In the schematic, all three species have a tandem array that is identical at the sequence level, but species-specific monomer start and end definitions lead to poor alignments downstream that are a necessary step in homology detection. As a result, this satDNA may be assumed to have relatively low similarity across species even though the array is identical at the sequence level. Early in the present research we encountered this issue commonly. (C) By making initial alignments, identifying a column that lacks missing data, cutting sequences, and swapping the left-hand sequence to the right end of its counterpart, monomers can be defined in a standard way across species and the satDNA can be correctly identified as having high sequence similarity following realignment. Our custom homology pipeline scripts employ the approach outlined in C to improve homology detection across *Plataphus* species.

**Figure 4.**
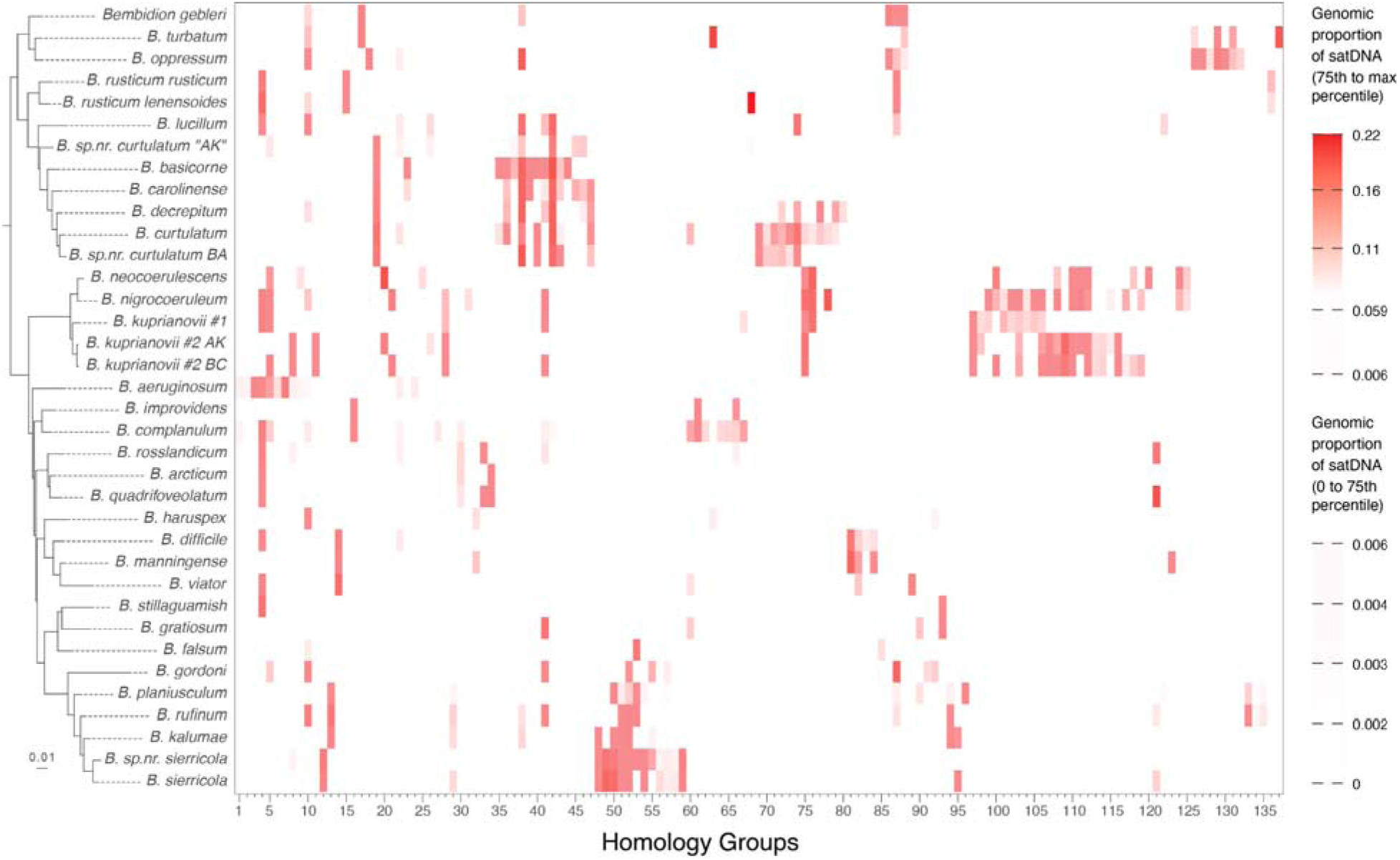
Turnover of homologous satDNA families across the subgenus *Plataphus*. Heatmap showing the genomic proportion of each satDNA family across species. Values are log-transformed to compress the wide range of proportions and improve visibility of low-abundance families. White cells indicate absence of a given satDNA family in a species’ genome. Two color bars are shown to visualize the full gradient: the lower bar spans 0 to the 75th percentile of nonzero values, while the upper bar spans the 75th percentile to the maximum. The midpoint of the color gradient is anchored to the median nonzero proportion across all detected families. *B. sp.nr. curtulatum BA* is referred to as *B. sp.nr. curtulatum “Bay Area”* in Figure 1.

Like subgenus-wide patterns, homology analysis in the *breve* species group showed evidence of limited satDNA conservation and major shifts in satDNA abundance across closely related species (Fig. 4). We detected 46 satDNA homology groups for which a putatively homologous satDNA was present in at least two different species. The average number of species per homology group was 2.9 and only eight satDNA homology groups had representation of at least five of the eight species in the group. Repeat profiles of six of the most broadly conserved satDNAs in the group showed some satDNAs were conserved with low to moderately low copy number while others show major shifts in copy number across species. Most notably, one satDNA (see satDNA 20, Fig. 5) showed low to moderate average copy number (e.g., >2K – 15K copies) in most species in the group; conversely, that satDNA was estimated to have >1 million and >250,000 copies in *Bembidion laxatum* and *B. ampliatum*, respectively. In the case of *B. laxatum*, that amounts to hundreds of megabases of sequence belonging to that one satDNA motif. This is more than a 2,000-fold increase in abundance compared to the estimated copy number of close relatives like *B. lividulum,* despite these species showing very little divergence in terms of external morphology and protein-coding gene sequence evolution [40].

**Figure 5.**
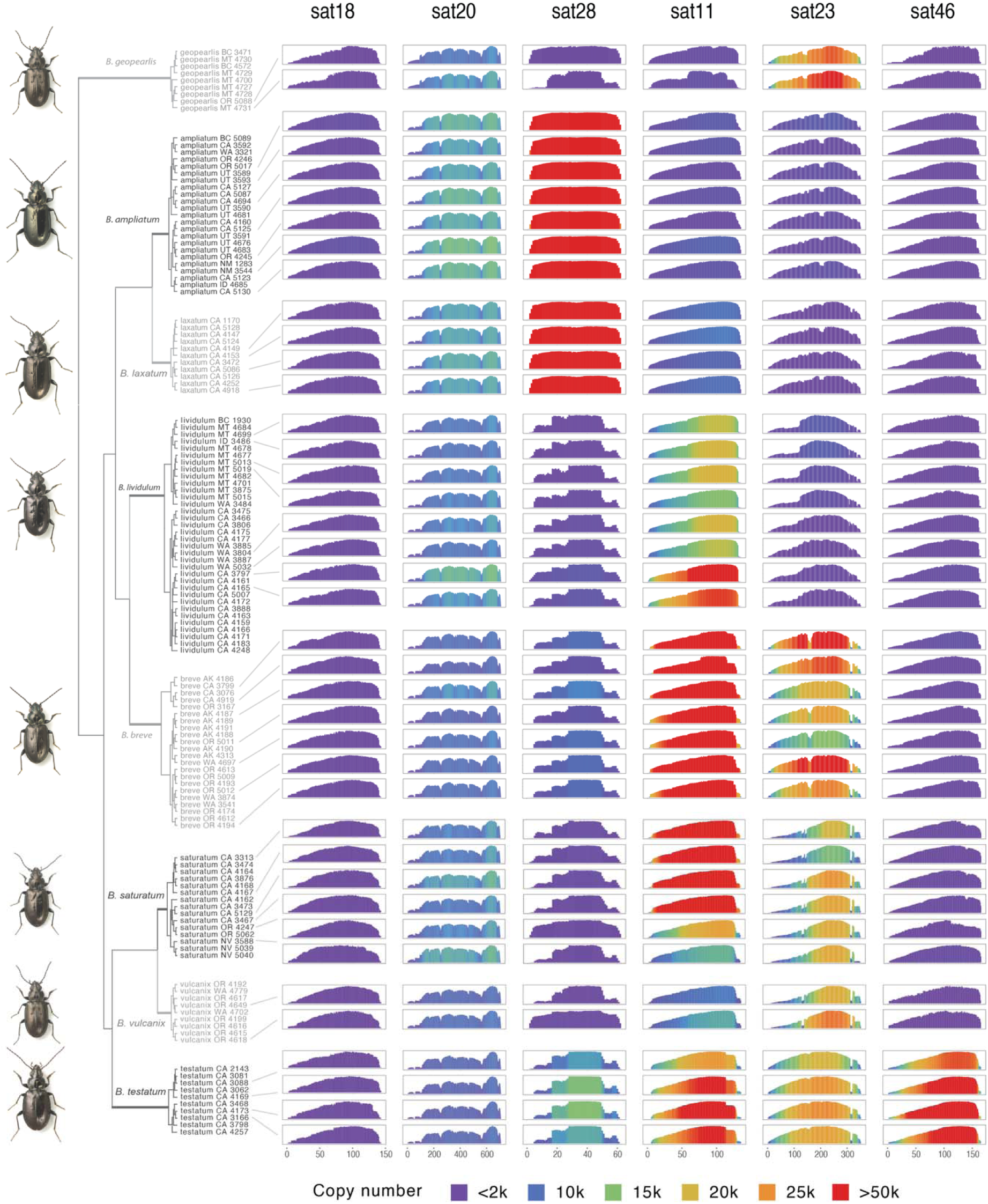
Shared satDNA profiles in the breve species group. Copy number profiles for six selected satDNAs shared across multiple individuals of the eight closely related species in the *breve* species group. Selected examples include satDNAs with uniformly low or moderate abundance across all species (i.e., 18 and 208), as well as satDNAs with variable abundance that ranges from relatively few copies in some species to over 1 million copies in other species (e.g., 28). Collectively, these account for tens of megabases of genomic variation among species (28, 11, 23, 46). SatDNA copy number estimates for those satellites shown range from less than 100 copies to more than 1.8 million.

Validation of our satDNA detection methods and homology pipeline with well-studied *Drosophila* species indicated overall robustness in our methods. RepeatExplorer2 identified all expected satDNAs (i.e., *Rsp*, *Rsp-like*, and *1.688* family satDNAs) in one or more target species.

Consistent with our satellite-specific predictions, *Rsp* was only detected in *D. melanogaster*, and thus was not present in any homology groups produced by our pipeline. *Rsp-like* was detected in all three predicted species (*D. mauritiana, D. simulans,* and *D. sechellia*) and our homology pipeline placed all three in the same homology group. For *1.688* family satellites, RepeatExplorer2 identified this target in all six predicted species (*D. melanogaster*, *D. simulans*, *D. mauritiana, D. sechellia, D. yakuba, and D. erecta*). Our pipeline grouped these satDNAs into two homology groups (one containing *D. simulans*, *D. melanogaster*, *D. mauritiana, D. sechellia*), and the other containing (*D. melanogaster, D. yakuba, and D. erecta*). The presence of two homology groups for *1.688* family satDNAs is consistent with the fact that monomers in this family are known to show a fairly old history of diversification and notable sequence-level diversity within and across species (e.g., much more than *Rsp-like*) [48,49]. In summary, these validation steps showed consistent RepeatExplorer2 detection of well-studied target satDNAs across *Drosophila* species and our homology pipeline sorted target satDNAs into groups consistent with expected patterns reported in the literature.

### Validating patterns with long-read data in *Bembidion laxatum*

Long-read PacBio sequencing for *B. laxatum* produced 29.84 Gb of output. Following assembly and removal of duplicate contigs and contaminants, the final assembly length was 1.42 Gb with a contig N50 of 3.7 Mbp, on par with other recently sequenced ground beetle genome assemblies. The genome assembly was highly complete as assessed with compleasm and the Endopterygota odb10 BUSCO set (99.2% complete–95.48% single copy, 3.72% duplicated).

Our primary goal in analyzing this assembly was to validate the striking abundance of a single satDNA motif which showed an average genomic abundance of 24.4% across all *B. laxatum* specimens sampled, with a maximum value of 32.2% in one specimen. When we used the consensus sequence of this abundant satDNA monomer (detected by RepeatExplorer2) as a query sequence in a BLAST search against the custom repeat library generated by EarlGrey, we observed hits from several sequences within the repeat library. Of these, three stood out as being highly abundant within the assembly – two of these were classified by EarlGrey as “satellite” and one was classified as “Unknown”. Each was composed of the query sequence (i.e., the consensus sequence of the abundant satDNA identified by RepeatExplorer2) tandemly arrayed in a few of its variant forms. Together, these three repeats alone account for nearly 422 Mb of sequence in *B. laxatum*, which is 29.7% of the assembly. This value falls toward the higher range of values estimated from multiple replicate *B. laxatum* individuals for which we report short-read estimates above.

In addition, we conducted BLAST searches of all 16 other consensus satDNA sequences identified from short-read data for this species. We found all 16 to be both present in many copies within the assembly and forming tandem arrays either composed of individual monomers or as monomers embedded within a higher-order tandem repeat structure, as is common with satDNA [8]. As such, our long-read validation strongly corroborates the high estimated abundance of the single-most abundant satDNA motif observed across all *Plataphus* study species. It also corroborates the general accuracy of satDNA detection in RepeatExplorer2 which we relied upon for the core short-read dataset.

### Satellite DNA abundance survey across insects

Our sampling more broadly across beetles and other insect orders showed that species in subgenus *Plataphus* are notable in both their genomic abundance of satellite DNA and variance in that value. Both mean satDNA proportion and variance were significantly higher in *Plataphus* than in every other lineage sampled (p < 0.05 for both Welch’s t-test and Levene’s test) (Fig. 6). However, we observed several cases in which individual species sampled from other lineages had similar or greater satDNA abundance than the average *Plataphus* species. Most notably, such species were present in the beetle suborder Adephaga, the beetle order Coleoptera, and the order Hymenoptera (bees, ants, and wasps) (Figs. 6, S4). Of these, Adephaga showed the highest mean satDNA abundance, and the highest variance, which were significantly higher than five of the eight other groups (i.e., all but Hymenoptera, Coleoptera, and *Plataphus*).

**Figure 6.**
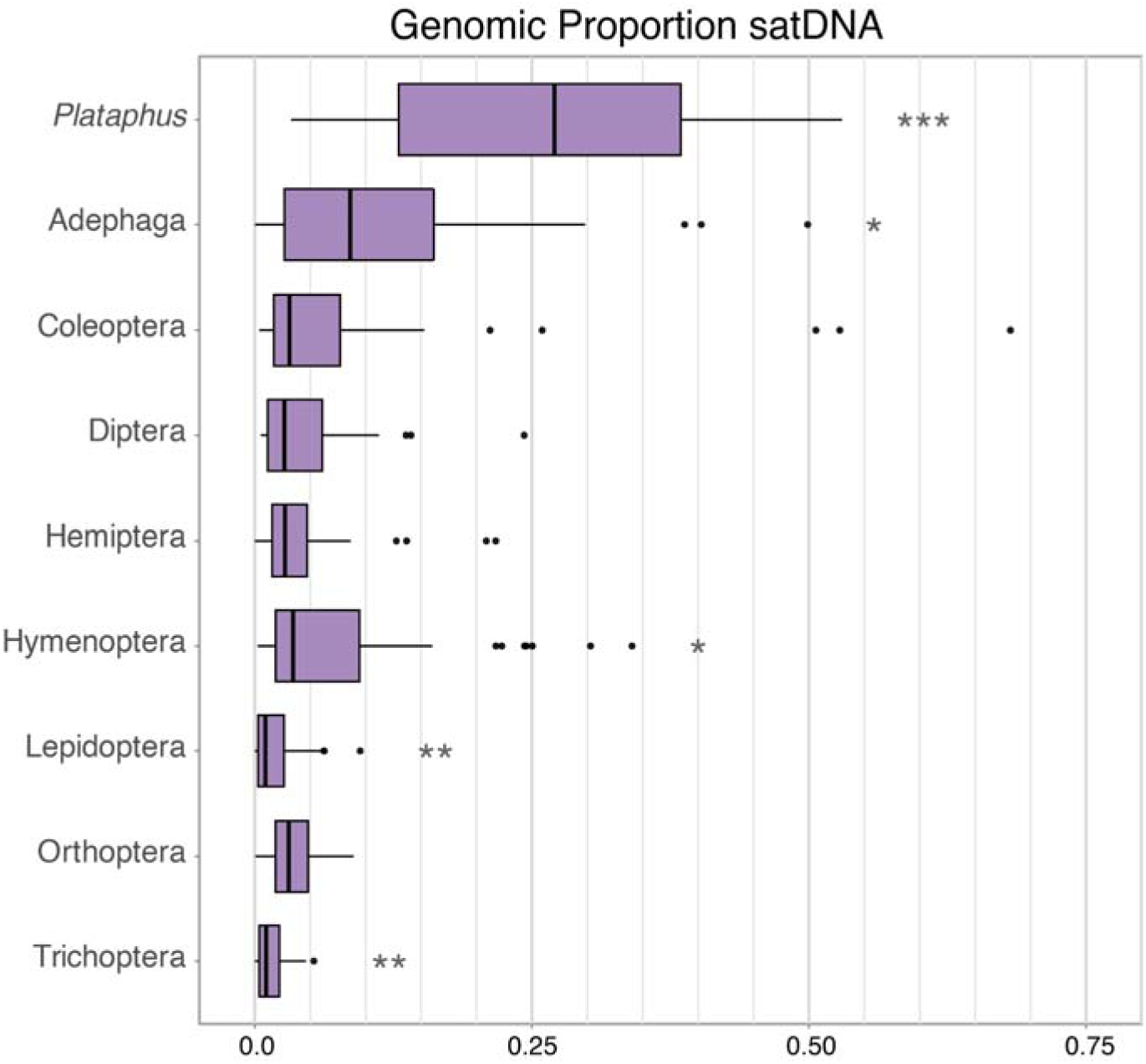
Analysis of satDNA genomic proportions across major insect lineages. Boxplots show satDNA genomic proportion (%) across *Plataphus*, Adephaga, Coleoptera, and the other six focal insect orders. Stars summarize Welch’s t-test results (BH-adjusted p < 0.05): *** = significantly different from 8/8 other lineages, ** = 7/8, * = 6/8. Variance-significance trends (Levene’s test; p < 0.05) are similar overall, except Lepidoptera variance is significantly different from 5/8 lineages rather than 7/8. Full pairwise p-values for Welch’s t-tests and Levene’s tests are reported in Tables S1 and S2.

Hymenoptera mean satDNA content and variance were significantly higher than most other lineages, as well (Fig. 6). Within both Adephaga and Coleoptera were species with satDNA genomic proportions that approached or exceeded 50% (Figs. 6, S4). In contrast, Lepidoptera and Trichoptera had notably low mean satDNA proportions, significantly lower than all other sampled lineages, though they did not differ significantly from one another (Fig. 6). In addition to group-level summaries in Figure 6, Supplemental Figure S4 reports satDNA proportions for all samples within each lineage and Supplemental Tables S1 and S2 report statistical comparisons of satDNA abundance and variance across species within lineages. Our sampling sensitivity analysis showed generally modest shifts in mean and variance values across focal groups when we increased sampling from 25 to 50 species within groups (Fig. S5).

## Discussion

Satellite DNAs (satDNAs) can be among the most dynamic components of eukaryotic genomes [7]. Here we characterize satDNA dynamics in a many-species framework of ground beetles, and place findings in a broader context of satDNA abundance estimates sampled across 400 insect species. Taken together, our investigation of satDNAs at both shallow and deep evolutionary scales represents the broadest taxonomic survey of satDNA of which we are aware and highlights ground beetles in the target subgenus and the suborder to which they belong as potentially powerful systems for study of satDNA evolution. We discuss our findings and their implications in more detail in the sections that follow.

### High satDNA abundance in *Plataphus* and other beetle groups

Examining satDNA dynamics across 50 *Plataphus* species revealed that overall satDNA abundance estimates vary substantially within the subgenus (Fig. 1), ranging from just 3.3% in *B. arcticum* to 53% in *B. gordoni*. Our observation of 12 species with abundance estimates ranging from 40–53% genome proportion of satDNAs places these species at the higher end of estimates for any species reported in the literature [5,63,64].

We placed abundance estimates for *Plataphus* in a broader evolutionary context by randomly sampling 400 insect species across beetles and other insect orders to show that the abundance of satDNA in the focal subgenus *Plataphus*, is significantly elevated compared to other groups (Fig. 6). However, our broad sampling also showed that the trends we observed in subgenus *Plataphus* may be part of a broader pattern as the beetle suborder Adephaga, the group containing ∼47,000 species [61] to which *Plataphus* belongs, also shows significantly elevated satDNA abundance compared to most other insect groups we investigated (Fig. 6). Given that our sampling of Adephaga did not include any species in the subgenus *Plataphus* (Fig. S4) sampling from the focal subgenus did not help drive this result for Adephaga.

Our detection of high counts (i.e., over 40) of distinct satDNAs motifs in several species also suggests notably high diversity of sequences in the library of satDNAs in a given species compared to many groups [30]. Interestingly, the count of satDNAs detected was not strongly correlated with overall satDNA genomic proportion (Fig. 2C). This was likely because in these beetles relatively few individual satellites tend to account most of the overall satDNA content within species. The single most abundant satDNA across species accounted for an average genomic proportion of 8.9% (Fig. 2A & B) and in the case of a *Bembidion laxatum* specimen, a single satDNA comprised 32.2% genomic proportion (Figs. 1C, S4), a finding we further validated with long-read data. These findings of overall abundant satDNA and disproportionate abundance of individual satDNA motifs corroborate previous cytogenetic and sequence-based work in the subgenus which demonstrated that individual rDNA-derived satDNA motifs can show striking abundance in select species [41]. Our broader investigation here suggests that striking abundance of individual satDNAs is the rule as opposed to the exception across the subgenus. A few other studies have reported individual satDNA motifs that account for large genomic fractions [31,65], including other striking examples within beetles [2,66]. These previous findings in beetles, together with our observation of high satDNA abundance in *Plataphus,* the beetle suborder Adephaga, and many outlier species with high abundance across non-adephagan beetles (e.g., *Mylabris sibirica* with a genomic abundance 68.2%; Figs. 6, S4) suggest that unusually abundant and dynamic satDNA evolution may be a broad characteristic of this species-rich order.

### SatDNA turnover on short evolutionary scales

In addition to observing high general satDNA abundance in *Plataphus*, our mapping of satDNA dynamics to the *Plataphus* phylogeny revealed major shifts in abundance among closely related species. Although notable shifts in sister taxa or close relatives have been reported for decades [65,67,68], our results add to only a few studies that have approached satDNA dynamics from a many-species framework. One such study generated satDNA abundance estimates of 37 species in the genus *Drosophila* using similar methods and noted dynamic variation that ranged from 0.54% to 38.8% genomic proportion in the genus [35]. However, the patterns we observe in *Plataphus* beetles somewhat contrast the results across *Drosophila* by showing more striking shifts within species groups. For example, satDNA abundance patterns in *Drosophila* showed distinct phylogenetic signal across the genus [35], whereas both tests of phylogenetic signal in the current study suggest satDNA abundance metrics are poorly correlated with phylogenetic relatedness in *Plataphus*. This underscores the dynamic turnover of satDNA in the target system, especially given that the study species have likely diverged over a shorter time frame than species in the genus *Drosophila* [43,69].

Our homology analysis and quantification of satDNA turnover in RepeatProfiler provides additional evidence of satDNA turnover rates in these species. Our finding that no satDNA homology groups showed high sequence-level conservation across the subgenus, and few were conserved across more than 20% of the 50 target species (Fig. 4), provides a well-sampled illustration of the ephemeral nature of satDNAs within lineages over evolutionary time. Recent divergence dating in bembidiines that included sampling of *Plataphus* specimens estimated that most *Plataphus* species have diversified in the last 10–20 million years [43]. Given those estimates, our results would indicate that overall satDNA content among species likely shows near complete turnover every 2–7 million years.

At finer taxonomic scales within species, we observed only modest variation within replicate individuals of the same species (Fig. 1C). This finding, including the results of our variance partition analysis, is consistent with theoretical expectations of concerted evolution in tandem repeats like satDNAs [28,70] and corroborates empirical observations of previous studies [71].

### Accelerating homology detection of satDNAs

A core obstacle to studying satDNAs in a many-species framework is that establishing homology of specific satDNAs can be a challenge. Even simple steps like producing high confidence alignments of putative homologs can present unintuitive technical challenges (Fig. 3). As we developed our homology pipeline, we noticed unexpectedly low levels of homology detection when analyzing consensus sequences generated from well-studied *Drosophila* species. Through further investigation, we realized that the TAREAN algorithm employed by RepeatExplorer2 may commonly define the start and end points of satDNA monomers differently in different species, even if a highly conserved satDNA is present in each species (see Fig. 3 for a schematic outlining this issue). This is not surprising or inherently problematic behavior – any start and endpoints defined in satDNA arrays are for human convenience rather than to detect biologically real start and end points. However, the species-specific nature of the output does present unintuitive challenges to downstream analysis which relies upon alignments to establish homology. To resolve this common occurrence, we added to our pipeline a step which standardizes start and endpoints of putative homologs through initial alignment to a dimerized repeat followed by end swapping and realignment. This and other experiences during this project underscored to us that despite the increasing availability of useful tools to detect and analyze satDNA, this is an area of genomics research that would benefit from increased emphasis on tool development [72], particularly to accelerate study in a multi-species framework.

### Implications for the library and centromere drive hypotheses

Key hypotheses that have framed much of the satDNA research in recent decades are the library hypothesis [65], and the centromere drive hypothesis [8,28]. These hypotheses seek to explain observed patterns of satDNA turnover which can be driven by stochastic processes such that species-specific patterns of abundance appear random (i.e., drift-like) [28,65,73].

Alternatively, abundance patterns can appear directional with certain satDNAs showing unexpectedly high proliferation consistent with selection (i.e., meiotic drive-like) [28]. Our data could conceivably fit both explanatory frameworks. The pattern of turnover evident in our analysis of homology groups (Fig. 4) indeed shows apparently stochastic expansion and loss of a subset of satDNAs which account for species-specific differentiation, with the library of satDNAs itself showing somewhat rapid turnover across evolutionary time. In addition, we report cases where individual motifs account for disproportionately large genomic fractions and show striking shifts (e.g., 2000-fold increases) in abundance between closely related species consistent with drive-like expectations (Figs. 1 & 4). Since Dover’s original proposal of drive-like evolution accounting for some satDNA dynamics, explanatory mechanisms centered in centromere evolution have found traction [8,15].

Determining whether the striking satDNA evolution in these beetles fits models of centromere drive would require extensive additional analysis including development of antibodies to determine whether satDNAs involved in recent, major satDNA proliferations are in fact centromeric in nature, and not just satDNAs embedded in peri-centromeric heterochromatin. However, our investigation in *Plataphus*, and broader survey of insect satDNA dynamics may help identify promising systems that could further test primary hypotheses of satDNA evolution.

### SatDNA abundance and genome architecture strategies in insects

Our broader comparison across major insect lineages showed interesting variation in satDNA genomic proportion among groups (Fig. 6). In addition to finding abundant satDNA in many beetle lineages, our data suggests that on average hymenopterans have relatively large amounts of satDNA in their genomes. This finding corroborates cytogenetic and genomic studies in bees which have reported high satDNA abundance in some genera [74,75], including wide variance between two species in the genus *Melipona* in which satDNA abundance is closely associated with overall heterochromatin abundance [75].

Within Lepidoptera and Trichoptera, by contrast, both showed distinctly low satDNA content and low variance in our initial sampling of those orders (Fig. 6). Generally low satDNA abundance in Lepidoptera has been noted in other studies [76], including several species with >1% genomic abundance [76–78] consistent with our observations. However, we note that two studies report satDNA expansions in lepidopteran species with unusually large genomes [79,80] in which satDNA genomic proportions are as high as 14.3% (which exceeds our highest observed estimate for a lepidopteran in the current study of 9.5%). Within Trichoptera our results corroborate and expand upon previous observations of low satDNA abundance in analysis of 17 caddisfly genome assemblies using both long-and short-read data [81]. This study observed average satDNA abundance across the order to be 1.5% and the species with the largest genomes (e.g., 1–1.3 Gb) ranged from 3.0–5.1%. Additional satDNA studies in these sister orders compared to other insect groups may help clarify whether satDNA abundance can be predicted by chromosome type (e.g., most lepidopteran and trichopteran species have holocentric chromosomes [82,83], though Trichoptera is less well-characterized [84]), genome size, heterochromatin abundance, and sex-specific achiasmy during meiosis.

Although our broader sampling was designed to maximize the diversity of genera and represent most major lineages (e.g., families) within the target insect orders, relative to the staggering diversity of insects, 400 species represent a very small preliminary sampling. However, seeing initial variation across orders that indicate elevated satDNA in some groups (e.g., Coleoptera, Hymenoptera) and notably reduced satDNA abundance in others (e.g., Lepidoptera, Trichoptera) merits further investigation. We hypothesize that if patterns indicated in this initial sampling, or other group-specific patterns emerge, these could indicate core differences in order-level genome/chromosome architecture strategies employed by these groups.

### Conclusions and future directions

In summary, we characterize patterns of rapid satellite DNA evolution in a many-species framework of ground beetles belonging to the subgenus *Plataphus*. Within these species we observe high overall satDNA abundance, high abundance of individual satDNA motifs, and striking abundance shifts among closely related species. By quantifying rates of satDNA turnover we provide a strong illustration of the ephemeral nature of these rapidly evolving sequences. Our survey of satDNA diversity across major insect groups places our findings in a much broader evolutionary context and identifies preliminary trends in satDNA abundance across several major insect lineages. We identify and purpose solutions for unintuitive technical challenges associated with studying satDNA in a many-species framework, and we validate several key findings using a combination of analysis of well-studied *Drosophila* species and long-read data generated in a particularly compelling study species, *Bembidion laxatum*.

Although our goals in this paper were unrelated to species delimitation, our methods and findings are of potential interest to researchers studying species boundaries or population structure. For example, the high variation between and relatively stable signatures within species observed in multiple analyses (Figs. 1 & 5) illustrate the potential for satDNA data to provide strong evidence of species boundaries on short evolutionary time scales. This approach adds further support to recent studies that emphasize the potential of repetitive sequences as a powerful source of signal to accelerate understanding of biodiversity [41,72,85–87].

Beyond helping to characterize existing biodiversity, satDNA studies across phylogenetic scales may accelerate understanding of the mechanisms that give rise to that diversity. In some species, major shifts in satellite DNA abundance and chromosomal location have been shown to contribute to speciation, as divergence in repeat content can promote reproductive isolation among lineages [22,23]. From a practical standpoint, biodiversity-scale surveys of satDNA may help more efficiently identify study systems that could prove useful in elucidating the drivers of reproductive isolation, and whether core genome architecture strategies (e.g., satDNA-rich chromosomes) can predict lineage-specific rates of speciation.

Although analysis of short-read data similar to the approach taken here can help avoid bias in estimates of tandem repeat genomic abundance that may collapse in an assembly-based approach [45], advances in long-read sequencing make transformative resolution of repetitive DNA increasingly accessible across lineages [72,88]. Leveraging nucleotide-scale resolution available in long-read assemblies continues to show powerful applications in satDNA research [39,48,66,89,90] and enable more comprehensive understanding of the fine-scale mechanisms that shape the evolution of these dynamic genomic regions.

## Acknowledgements

We thank Dr. Beatriz Navarro Domínguez for sharing helpful satDNA data analysis experience and Uyiosa Osa-Iduma for help with data processing. We are indebted to Drs. Jiří Macas and Petr Novák for their generous resource accommodations and technical support during our use of the Galaxy RepeatExplorer2 portal. We are grateful to Professor Diogo Cavalcanti Cabral-de-Mello for reaching out with collegial feedback on our manuscript which improved the scope of our discussion of satDNA patterns across insects.

## Author Contributions

JSS conceived of the study and JSS, ZSW, SN, and AML contributed to aspects of the study design. ZSW, SN, JSS, PW, TP, LZ, GG, and PBF analyzed the data. SN and JSS led development of the novel homology pipeline. JSS, ZSW, SN, PBF and AML contributed to data interpretation. ZSW and JSS generated figures and wrote the first draft of the manuscript and all authors contributed to its subsequent refinement. JSS, AML, and ZSW contributed to funding the research.

## Data Accessibility and Benefits-Sharing

### Data Accessibility

All data files and code for analysis and producing plots are deposited at GitHub: https://github.com/johnssproul/satDNA_in_Plataphus. Output summary files are available in FigShare [DOI will be released upon publication]. NCBI accessions for newly generated long-read data in *B. laxatum* are deposited to NCBI Sequence Read Archive [BioProject: Accession will be released upon publication].

### Benefits Sharing

The authors collected the biological samples reporting newly generated data in this study. No genetic resources were obtained through third-party access-and-benefit-sharing agreements, and no traditional knowledge associated with genetic resources was utilized. The authors are not aware of any benefit-sharing obligations applicable to this work.

