## Supplemental Material for "Satellite DNA dynamics across phylogenetic scales in ground beetles and other insects"

Supplementary Methods

Survey of satDNA dynamics across major insect orders

We conducted additional analyses to compare patterns of satDNA content observed in the subgenus *Plataphus* of *Bembidion* (Carabidae) to patterns across much broader evolutionary scales by sampling 400 additional insect species. We moved outward from the subgenus *Plataphus*, to the suborder Adephaga (Coleoptera) with emphasis on the large family Carabidae to which *Plataphus* belongs, then to the order Coleoptera, and compared those patterns to the six most species-rich insect orders outside of Coleoptera (*i.e*., Diptera, Hemiptera, Lepidoptera, Orthoptera, Trichoptera, and Hymenoptera). This framework allows patterns observed among closely related species within *Plataphus* to be evaluated across increasingly broader evolutionary scales. Across all data sets, we standardized read depth to <1x genomic coverage. We then estimated satDNA abundance using RepeatExplorer2’s TAREAN module [1,2].

To generate the dataset we used for these 400 additional insect species, we assembled publicly available short-read genomic data. Prior to downloading data, we gathered metadata for focal clades, heavily filtered the initial list to reduce variables associated with use of public data, and then randomly sampled from the filtered list of candidate species within clades. Additional details on these steps follow below.

We used the Sequence Read Archive (SRA) Run Selector tool to download a run metadata for all species within a given order which met our initial filtering criteria. Initial criteria accepted only species for which data generation used the same sequencing platform (i.e., Illumina) and similar sample preparation and sequencing strategies (i.e., single-individual, 150 PE libraries, whole-genome shotgun sequencing). We then visually inspected the resulting metadata which included tens of thousands of samples. Thorough visual inspection helped us identify patterns that led to two actions. First, we noted a few categories of samples and study types that we subsequently excluded. These included cases where samples that actually represented pooled individuals, as opposed to single individuals, were not correctly filtered out at the SRA Run Selector step, and cases where data arose from microbial studies (especially metagenomics often focused on insect gut contents) which we subsequently excluded. Second, we identified genera for which a disproportionately high number of species were represented in public data relative to the overall diversity in the order to which they belonged. This led us to exclude the fly genera *Drosophila*, *Aedes*, and *Anopheles* which have been the focus of extensive research in classic model systems as records for these three genera account for >33% of all data entries on our initial filtered list but they make up a relatively small fraction of fly diversity overall.

From this initial candidate list we then used a custom script to randomly sample 60 species from each order with the condition that no more than one species per genus could be selected. (Our sampling of 60 species per order here exceeded our target of 50 species in anticipation of some samples being potentially excluded following post-analysis filtering which is described below). Because taxonomic bias is expected in public data we evaluated whether our random sampling of genera resulted in representation of major (i.e., species-rich) families within each order. Major families were identified using the Catalogue of Life Checklist [3]. This sanity check revealed one major sampling gap in the order Coleoptera in which by far the largest beetle family, Curculionidae, was omitted in initial sampling. We repeated random sampling of Coleoptera but with forced sampling of multiple Curculionidae species. In the remaining orders sampled we observed that random sampling of genera led to representation of many families (range = 10–23 families represented per order) including at least the three most species-rich families of each order.

We used RepeatExplorer2’s TAREAN module [2] to estimate satDNA abundance for each species randomly sampled within each order. TAREAN uses an assembly-free, graph-based clustering approach to identify repetitive sequences from short-read data. Because TAREAN is designed for low-coverage inputs (<1x genome equivalent), we downsampled reads to ≤1x genomic coverage prior to analysis. Specifically, we downloaded the raw FASTQ files for each species selected by our random sampling of filtered SRA taxa above and processed them with a custom Snakemake pipeline which trimmed adapters and low-quality bases using Trim Galore (Krueger et al., 2015) then downsampled reads to <1x coverage using seqtk (available from [[4]](https://github.com/lh3/seqtk)). Because most of our sampled species lack published genome-size estimates, we used NCBI to identify the lower bound of genome size within an order based on the assembly length of the species with the smallest chromosome-scale assembly. We then calculated the 1x read count for that genome, and downsampled reads to that value for all species in the respective insect order. This conservative approach ensured that sampling our analyses were likely to conform to RepeatExplorer2 ≤1x genomic coverage in the absence of genome size estimates for all samples. A consequence of this approach is that some samples (i.e., those with the largest genomes) have much lower coverage (e.g., 1/10 coverage) data analyzed in RepeatExplorer2 than the samples with the smallest genomes. This could lead to lower detection of satDNAs with low genomic abundance in species with large genomes. However, we preferred this loss in sensitivity of low-abundance satDNAs rather than risk overestimation of satDNA content that could occur if samples exceed 1x coverage.

Following analysis in RepeatExplorer2 we conducted post-analysis filtering of our data by visualizing summary graphs for all 480 candidate samples for which we conducted initial analysis to identify any samples with extreme trends of repeat abundance that could merit further scrutiny. During this, we identified a few samples that produced non-typical clustering results in which more than 90% of reads were assigned to repeat clusters in the RepeatExplorer clustering summary. For these samples, we examined SRA metadata and associated publications (when available) to determine whether the data were appropriate for inclusion in our analysis. Through this method, we identified and removed 22 runs derived from museum specimens (mostly part of the same study), mitochondrial genome sequencing projects, or non-focal lineages comprised of gut bacteria for which metadata led to conflated species identification and was not caught by our pre-analysis filtering. Museum-derived samples were removed because they can overestimate repetitive DNA content due to DNA degradation and associated sequencing biases.

One other sample raised questions during our visual inspection of RepeatExplorer output. The beetle species, *Mylabris sibirica*, had an estimated satDNA genomic proportion of 68.2%. This estimate was notably higher than the previous highest estimate of satDNA abundance for any species in the literature of which we were aware. We took two approaches to validate the estimates obtained from RepeatExplorer2. First, we tested whether this unexpectedly high result could be explained by coverage depth. This species was one of a few beetles in our sampling for which a genome size estimate was available as part of a whole-genome sequencing project [4]. Because it fell near the lower end of genome size, it was a sample for which reads input into RepeatExplorer2 were below but fairly close to 1X coverage – near the maximum recommended by RepeatExplorer2. We thus performed two additional RepeatExplorer2 runs for this sample using increasingly lower numbers of input reads (approximately 0.5x and 0.25x genomic coverage). Both of these subsequent runs returned similarly high satDNA estimates. We further validated this pattern by BLASTing the reported most abundant satDNA family against a published long-read assembly generated from the same data set. This approach made clear that the target satDNA is strikingly abundant across most scaffolds in the assembly [4]. As these further investigations corroborated our initial findings, we retained this sample and report its original results.

Following the pre- and post-analysis filtering steps described above to standardize our public data-derived dataset we conducted final randomized sampling from the remaining to obtain 50 samples per order (i.e., our target final data set of 400 species) using a Python subsampling script that evaluated 50,000 random candidate subsets and retained the subset whose satellite DNA proportion best matched the order-wide mean before reducing sampling to 50 species per order, which represents our final curated data set.

Supplementary Tables

**Table S1. Pairwise comparisons of satDNA genomic proportion among focal lineages.** Two-sided Welch’s t-tests were used to evaluate differences in mean satDNA genomic proportion between each pair of lineages. Values shown are Benjamini–Hochberg (BH) adjusted p-values. Bolded values indicate significant comparisons (p < 0.05).

|  | Plat. | Adep. | Cole. | Dipt. | Hemip. | Hymen. | Lepid. | Orthop. |
| --- | --- | --- | --- | --- | --- | --- | --- | --- |
| *Plataphus* |  |  |  |  |  |  |  |  |
| Adephaga | **<0.001** |  |  |  |  |  |  |  |
| Coleoptera | **<0.001** | 0.183 |  |  |  |  |  |  |
| Diptera | **<0.001** | **<0.001** | 0.072 |  |  |  |  |  |
| Hemiptera | **<0.001** | **<0.001** | 0.065 | 0.901 |  |  |  |  |
| Hymenoptera | **<0.001** | 0.052 | 0.817 | **0.025** | **0.020** |  |  |  |
| Lepidoptera | **<0.001** | **<0.001** | **0.004** | **0.002** | **0.004** | **<0.001** |  |  |
| Orthoptera | **<0.001** | **<0.001** | **0.030** | 0.357 | 0.430 | **0.004** | **<0.001** |  |
| Trichoptera | **<0.001** | **<0.001** | **0.002** | **<0.001** | **<0.001** | **<0.001** | 0.218 | **<0.001** |

**Table S2. Pairwise comparisons of variance in satDNA genomic proportion among focal lineages.** Levene’s tests (median-centered) were used to evaluate differences in variance between each pair of lineages. Values shown are Benjamini–Hochberg (BH) adjusted p-values. Bolded values indicate significant comparisons (p < 0.05).

|  | Plat. | Adep. | Cole. | Dipt. | Hemi. | Hyme. | Lepi. | Ortho. |
| --- | --- | --- | --- | --- | --- | --- | --- | --- |
| *Plataphus* |  |  |  |  |  |  |  |  |
| Adephaga | **0.010** |  |  |  |  |  |  |  |
| Coleoptera | **0.008** | 0.444 |  |  |  |  |  |  |
| Diptera | **<0.001** | **<0.001** | 0.083 |  |  |  |  |  |
| Hemiptera | **<0.001** | **<0.001** | 0.065 | 0.728 |  |  |  |  |
| Hymenoptera | **<0.001** | 0.115 | 0.713 | **0.035** | **0.022** |  |  |  |
| Lepidoptera | **<0.001** | **<0.001** | **0.016** | **0.034** | 0.083 | **<0.001** |  |  |
| Orthoptera | **<0.001** | **<0.001** | **0.021** | 0.065 | 0.154 | **0.002** | 0.472 |  |
| Trichoptera | **<0.001** | **<0.001** | **0.008** | **0.002** | **0.008** | **<0.001** | 0.083 | **0.002** |

Supplementary Figures

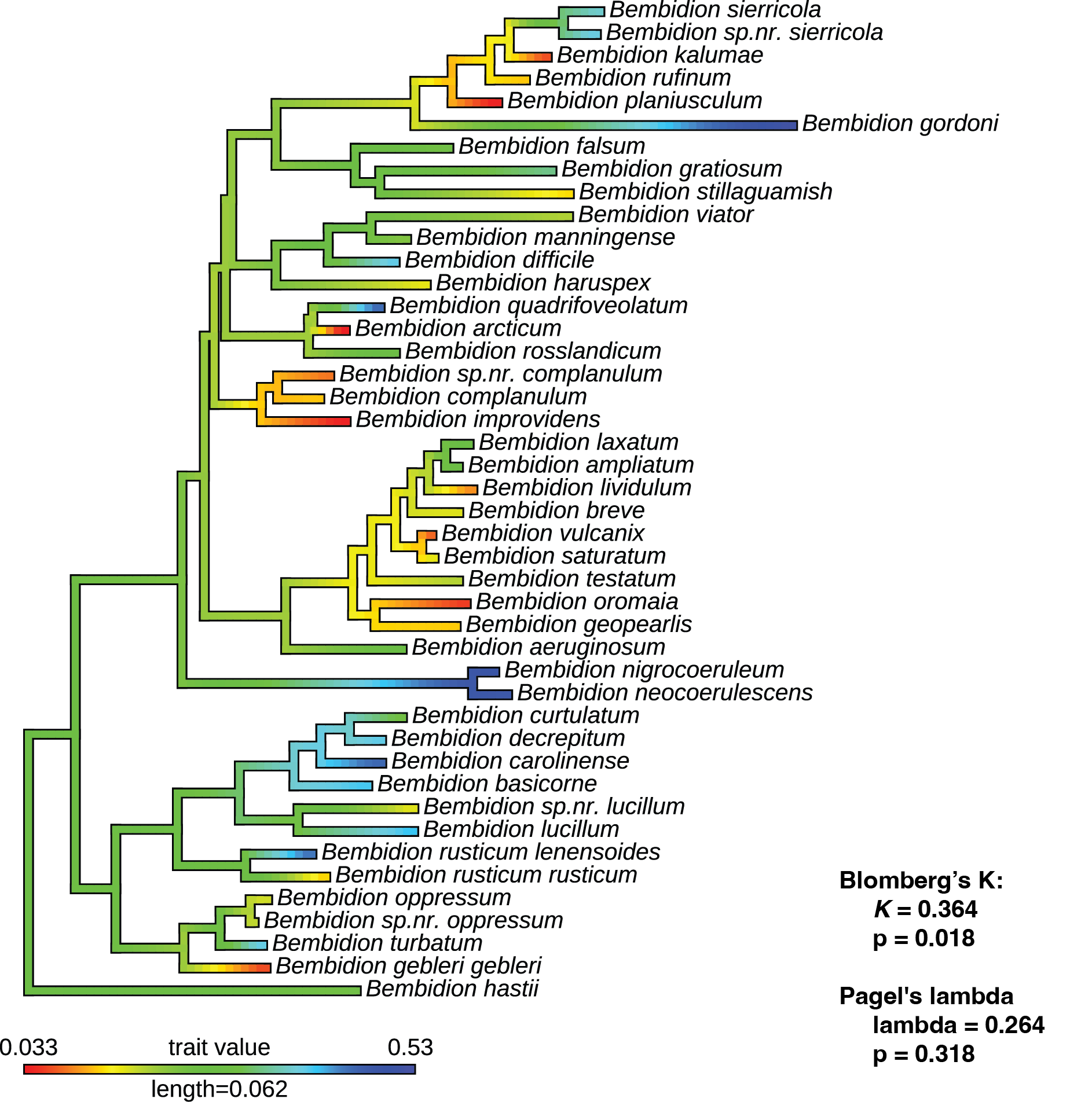

**Figure S1. Summary of phylogenetic signal tests for satellite DNA genomic proportion.** Tests of Blomberg’s *K* and Pagel’s lambda generated using the APE [5] and phytools 2.0 [6] packages in R [7].

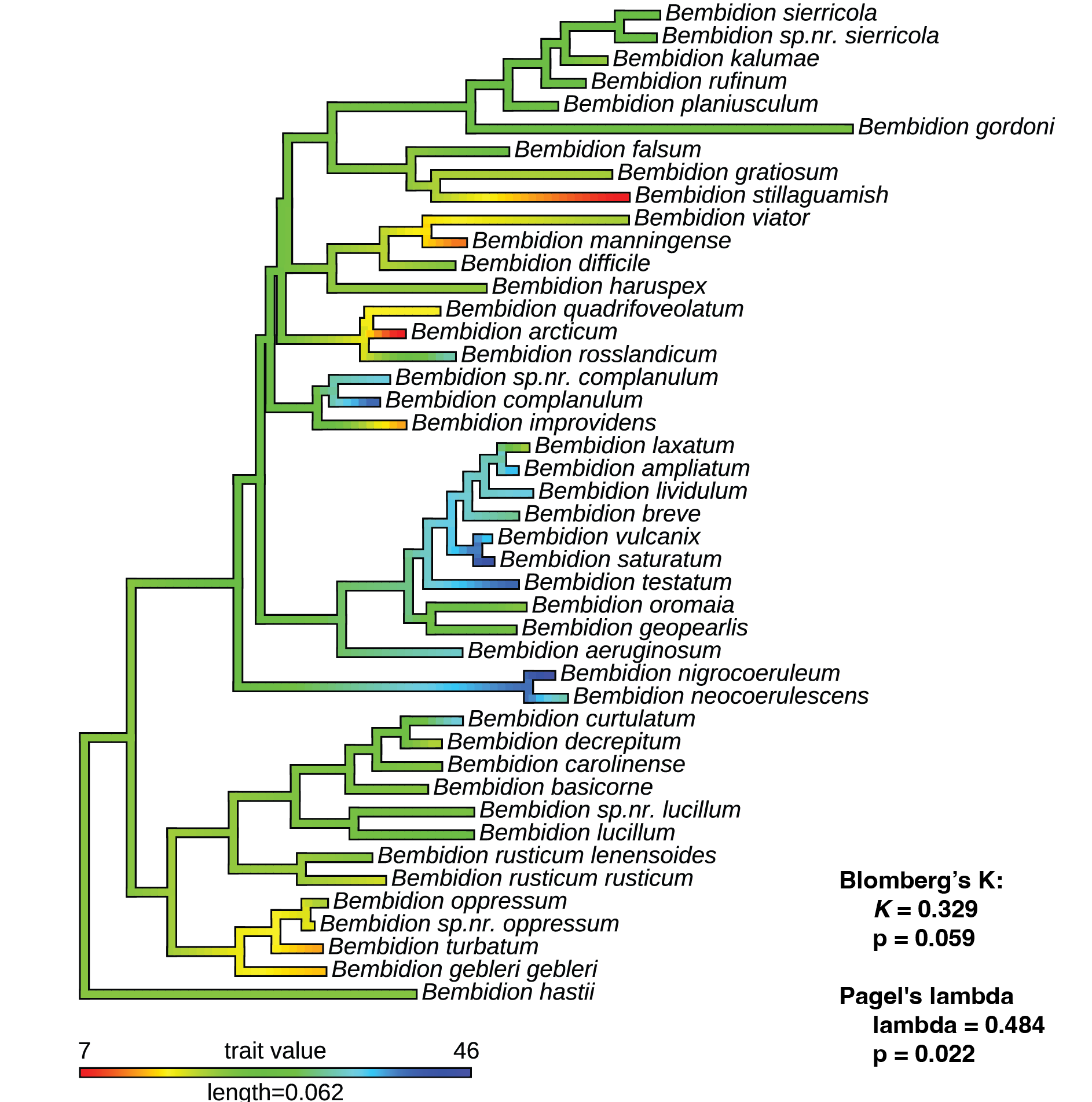

**Figure S2. Summary of phylogenetic signal tests for count of total satellite DNAs.** Tests of Blomberg’s *K* and Pagel’s lambda generated using the APE [5] and phytools 2.0 [6] packages in R [7].

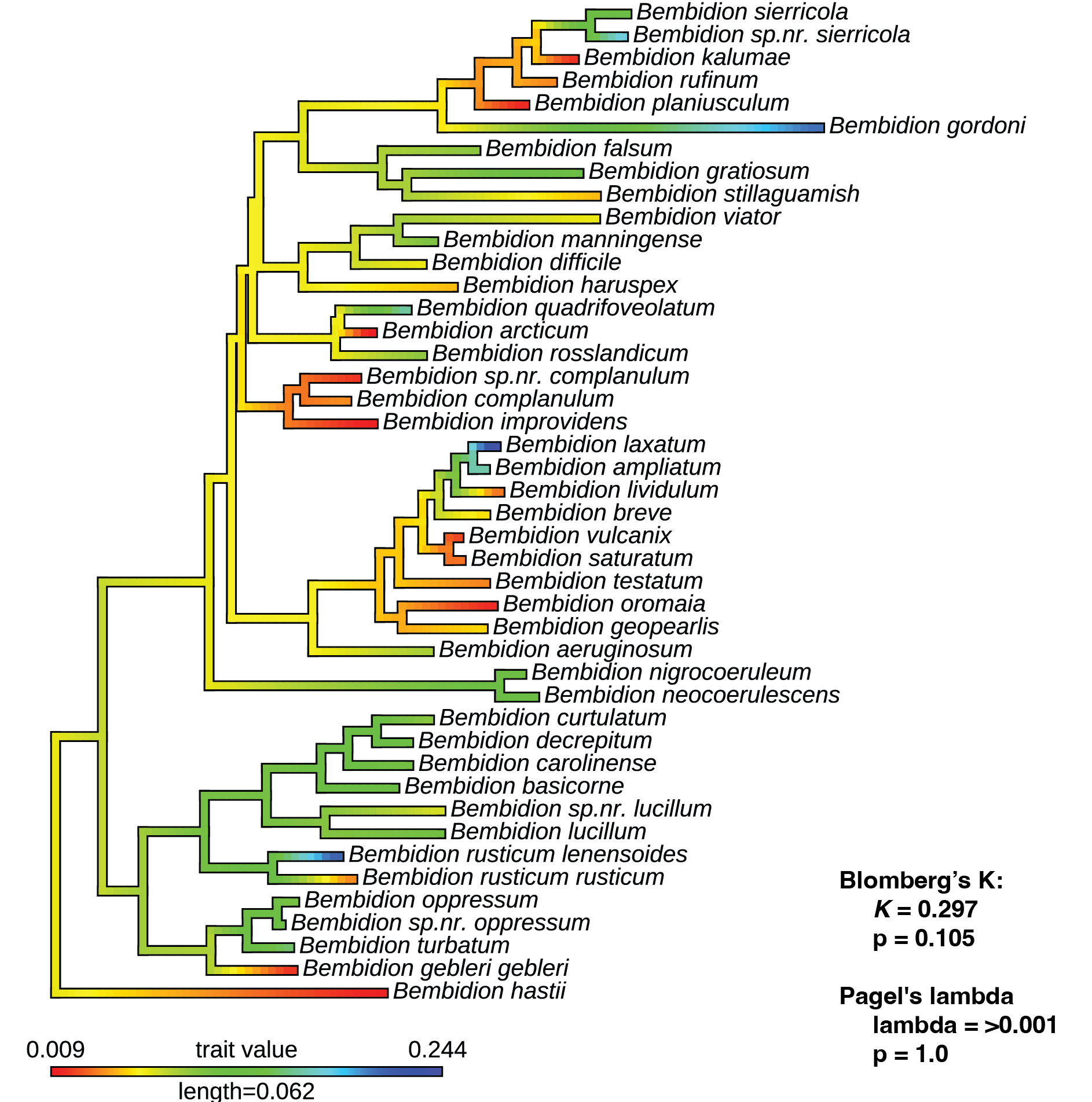

**Figure S3. Summary of phylogenetic signal tests for the single most abundant satellite DNA in terms of genomic proportion.** Tests of Blomberg’s *K* and Pagel’s lambda generated using the APE [5] and phytools 2.0 [6] packages in R [7].

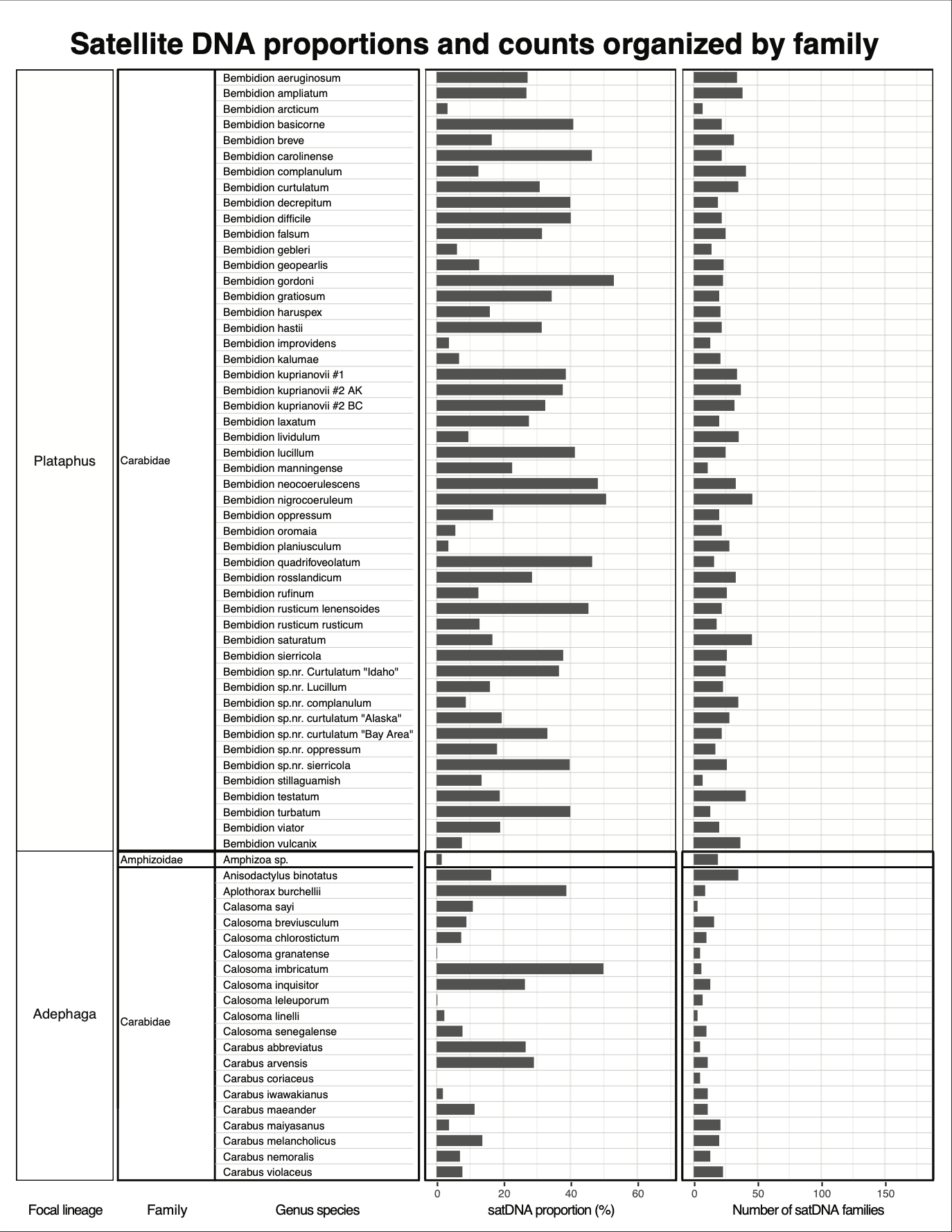

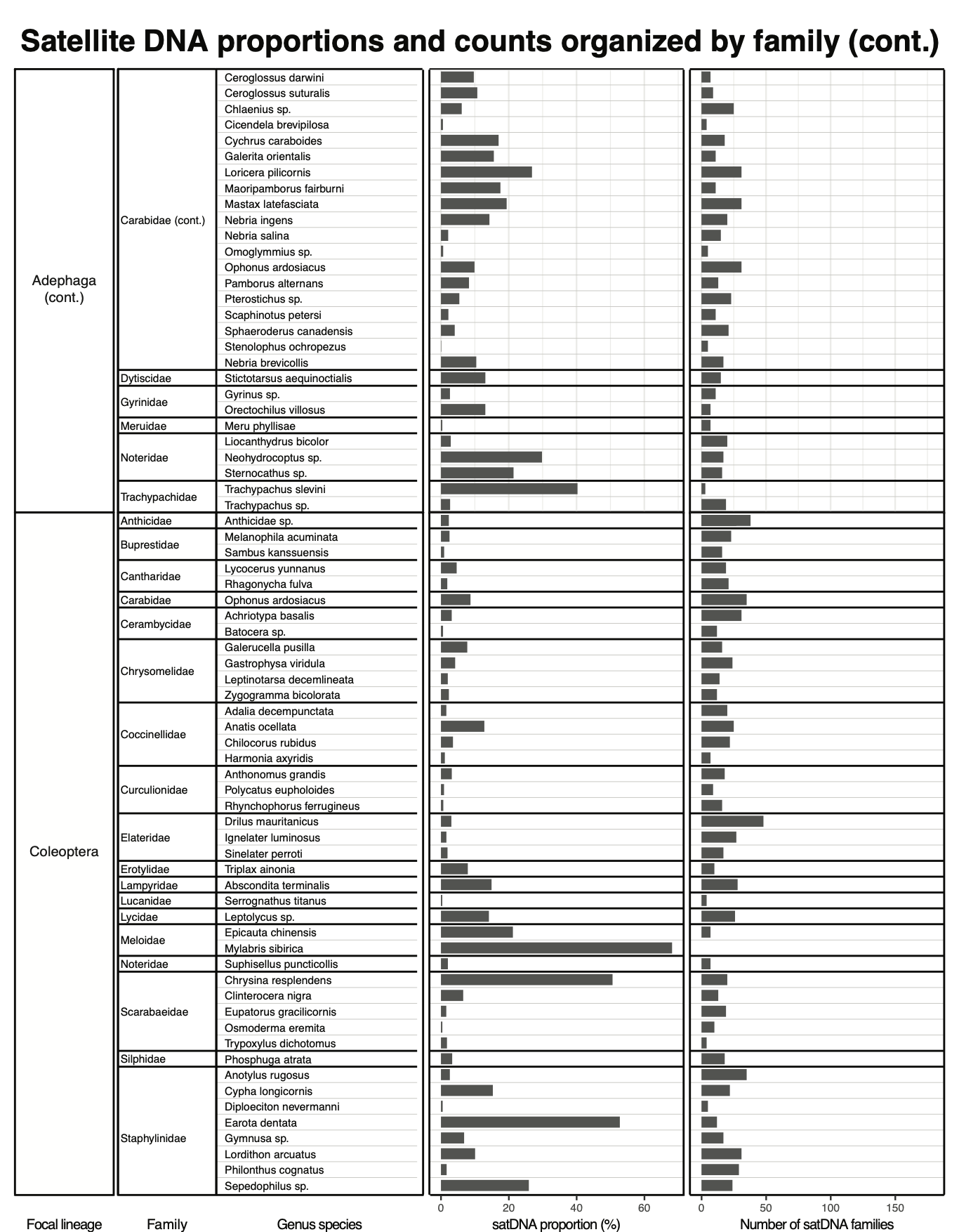

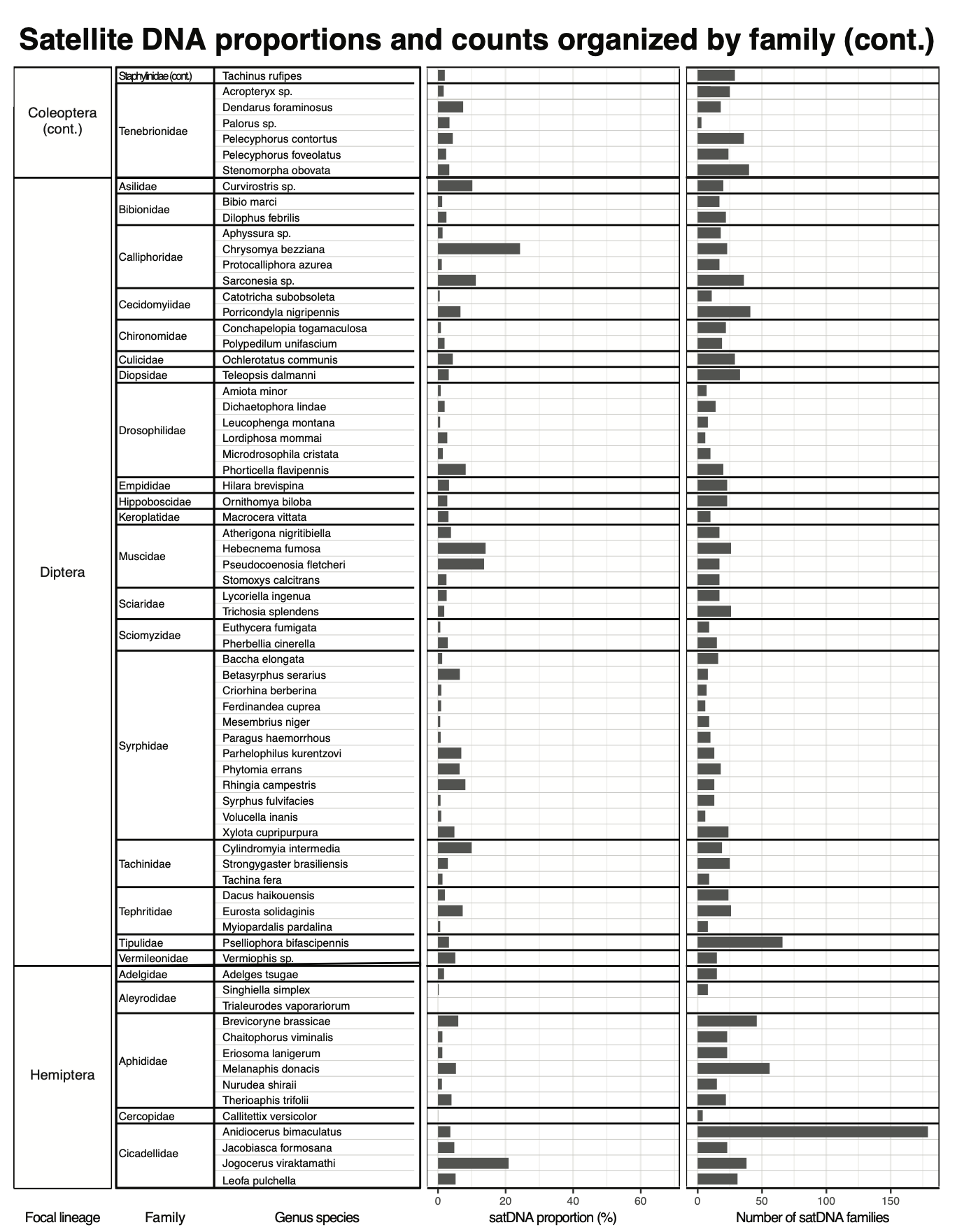

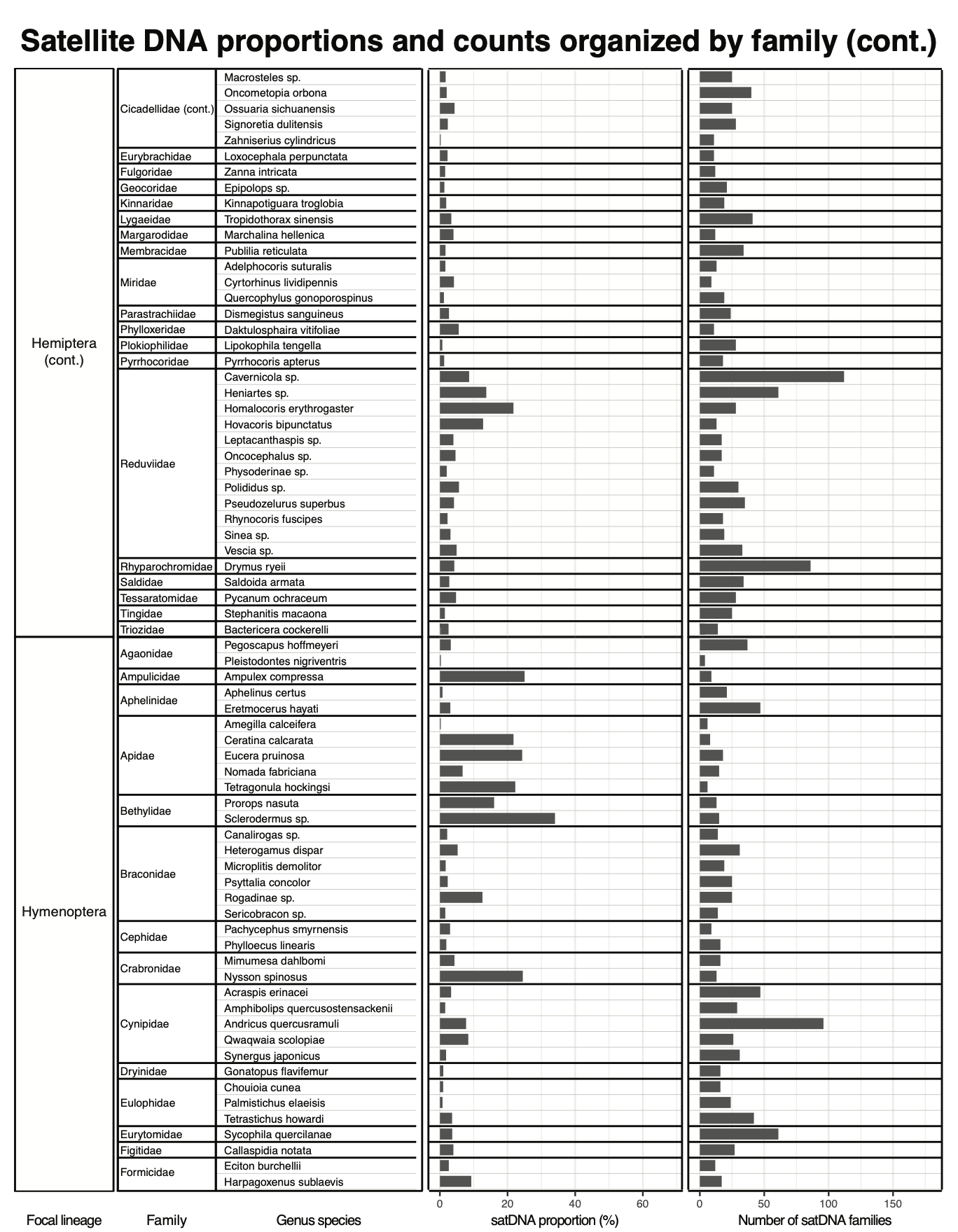

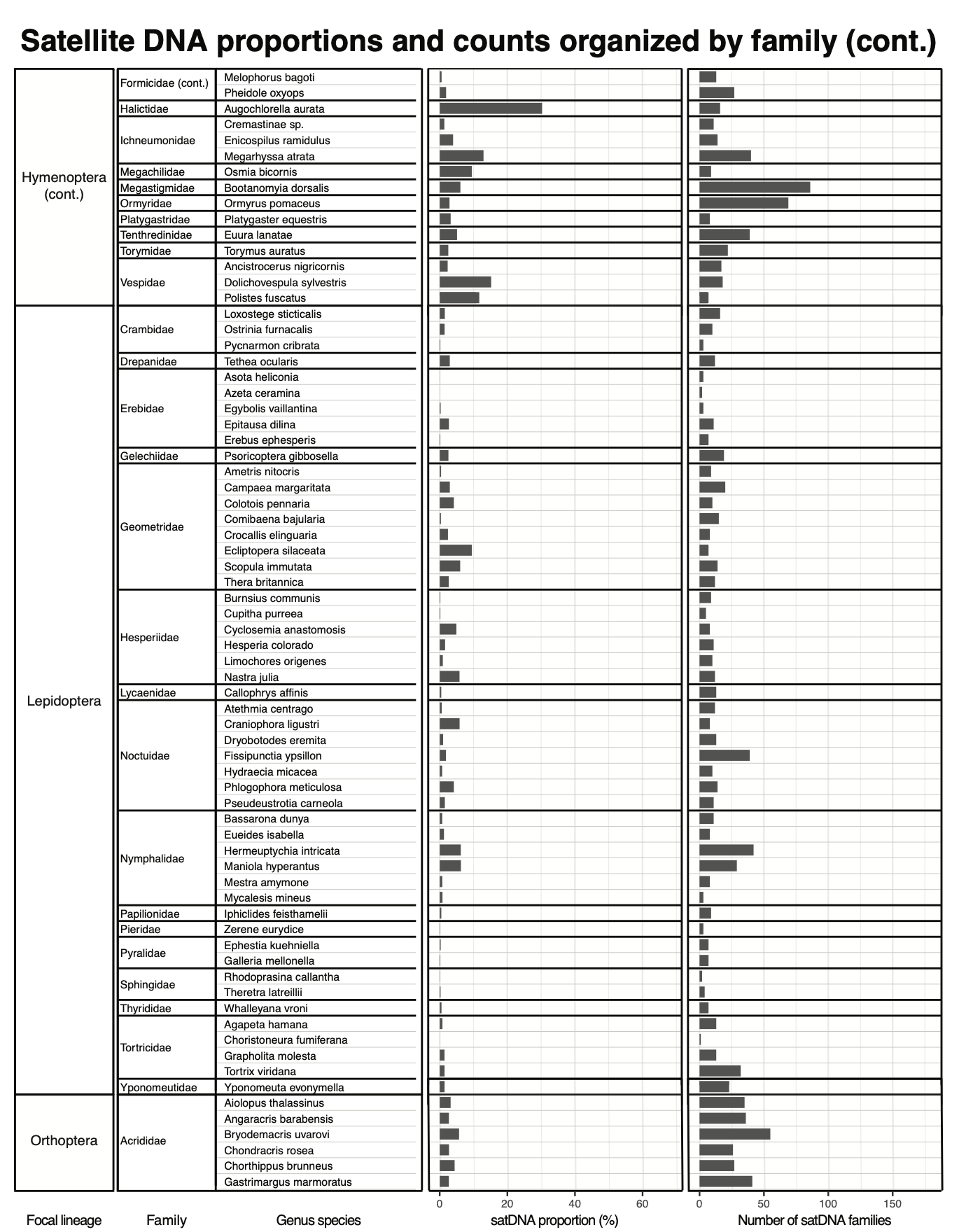

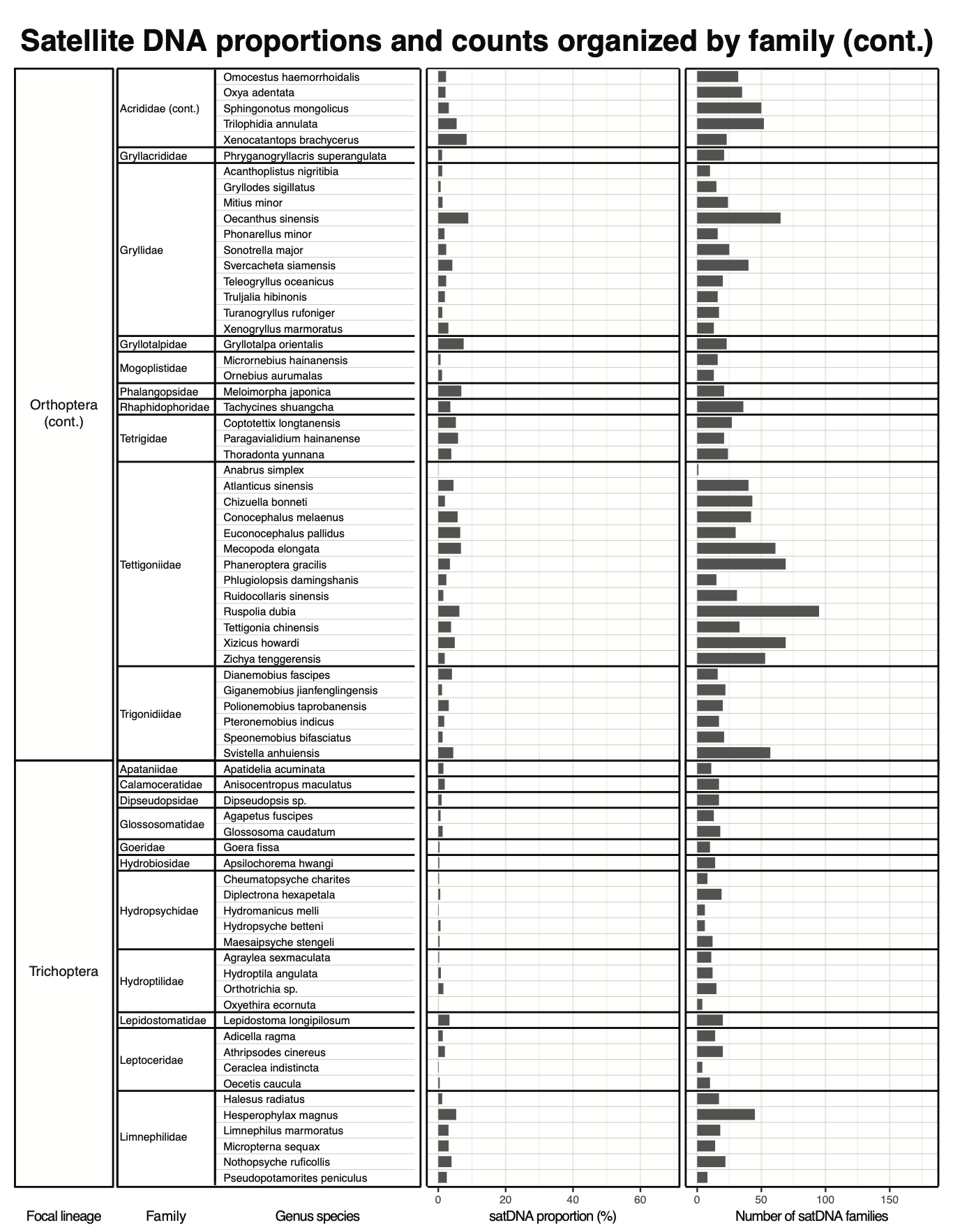

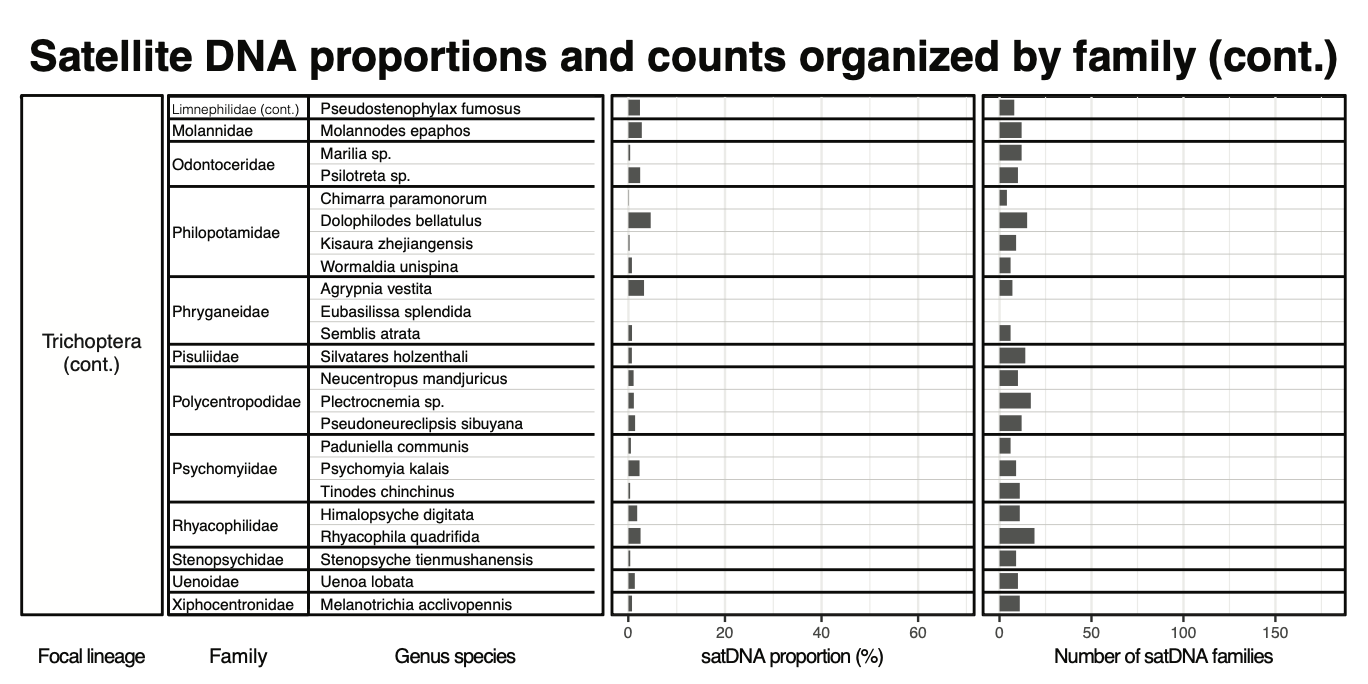

**Figure S4. SatDNA dynamics across all 450 focal samples, organized by taxonomic family.** Individual bar plots represent each specimen, illustrating their respective satDNA genomic proportion (%) and the total number of unique satDNA families identified within the genome. Samples are organized alphabetically within family groups.

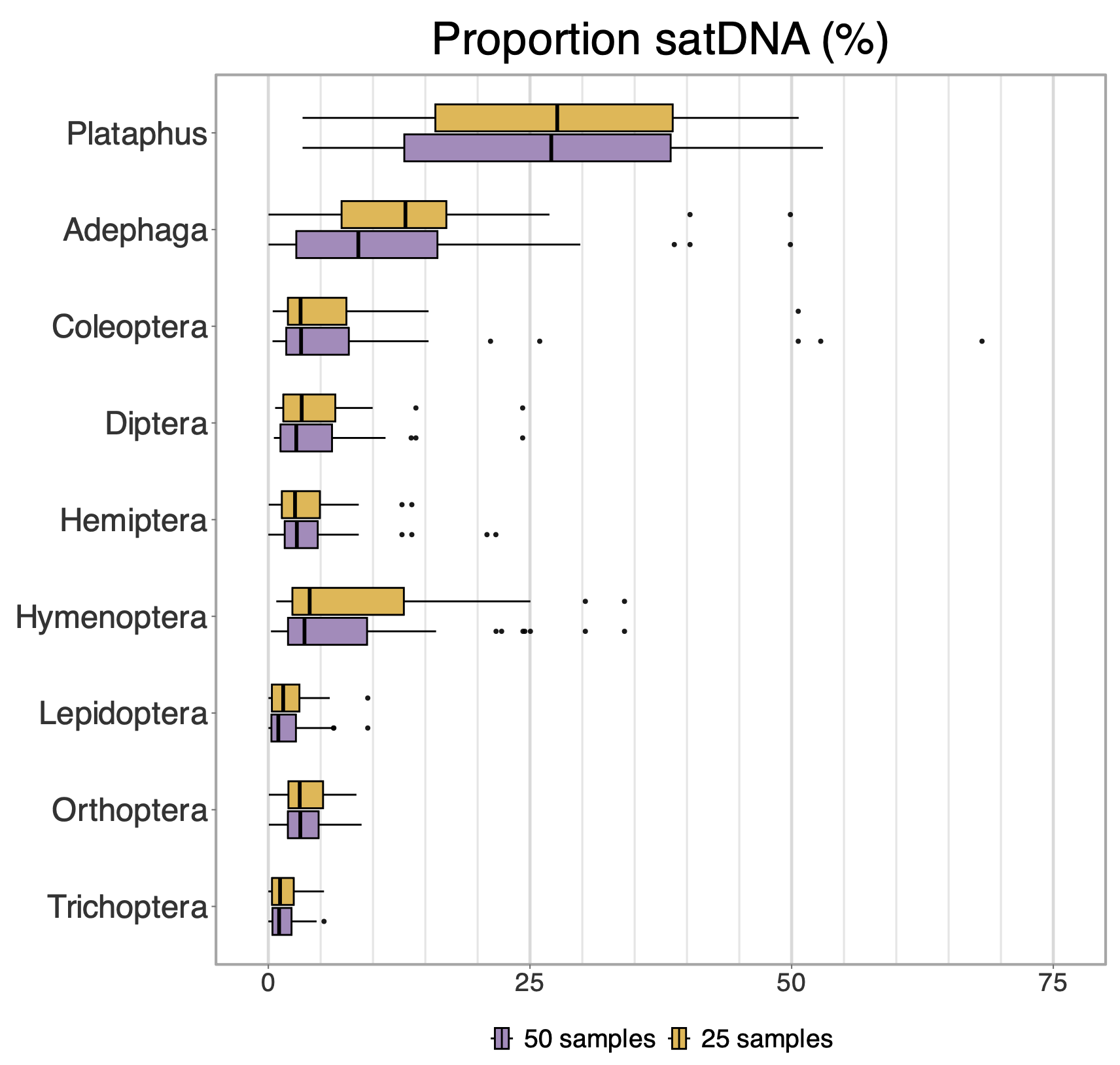

**Figure S5. Sensitivity analysis of satDNA genomic proportions across focal insect lineages.** Boxplots compare the distribution of satDNA genomic proportions (%) across the nine focal insect lineages using two different sampling scales: the full dataset of 50 samples per lineage (purple) and a random subsampling of 25 specimens per lineage (yellow). Individual outliers for both treatments are indicated by black points

7.	R Core Team. 2014 R: A language and environment for statistical computing.
